# Design of Fluorescent Membrane Scaffold Proteins for Nanodiscs

**DOI:** 10.64898/2026.04.07.716332

**Authors:** Estevan Cleveland, Aiden R. Wolf, Shawn Chen, Farhana Afrin Mohona, Isaac Kailat, Brian H. Tran, Suresh Babu Lavanya, Nicholas Primanis-Erickson, Yi-Chih Lin, Michael T. Marty

**Affiliations:** Department of Chemistry, University of Texas at Austin, Austin, TX, 78712, USA; Department of Chemistry and Biochemistry, University of Arizona, Tucson, AZ, 85721, USA

## Abstract

Nanodiscs are nanoscale lipid bilayer membrane mimetics surrounded by two membrane scaffold proteins (MSP). They are widely used as soluble cassettes for membrane proteins and lipids in diverse applications in structural, functional, and biophysical studies. The original MSP was derived directly from human apolipoprotein A-1, and novel constructs have been adapted from this original design, including fluorescent nanodiscs of varying designs. However, chemical derivatization with fluorophores can be expensive, and prior designs of split fluorescent proteins fused to MSP were limited in the color pallet available. Here, we developed MSPs with a wide range of different fluorescent C-terminal protein tags, including a versatile HaloTag fusion. These fluorescent MSP were purified following typical MSP purification procedures with similar yield. Then, we demonstrate that fluorescent MSPs form nanodiscs with similar structure and stoichiometry to conventional MSP nanodiscs. They are also suitable for assembly of nanodiscs with embedded integral membrane protein. Finally, we apply these constructs to monitor peripheral membrane protein binding to lipids via fluorescence resonance energy transfer. These fluorescent MSP constructs enable a range of different applications and provide a versatile template for future design of nanodiscs with unique functions.

**For Table of Contents Only:** 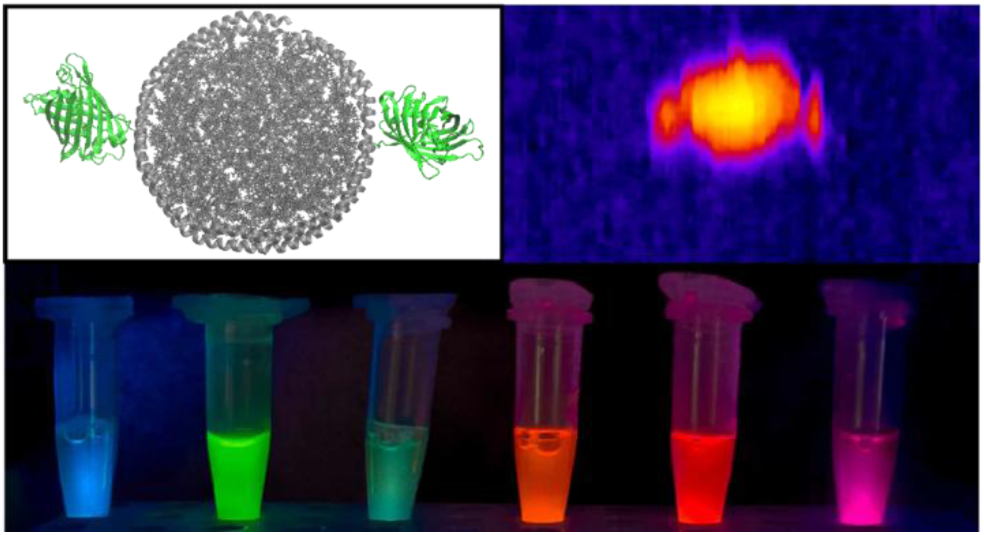

## Introduction

Since their initial release in 2002,^1^ nanodiscs have become widely used to study membrane proteins,^2–5^ protein-lipid interactions,^6–8^ and membrane biophysics.^9–12^ They have become especially useful for structural biology applications in cryo-EM.^13,14^ Nanodiscs contain a nanoscale lipid bilayer that is encircled by two amphipathic membrane scaffold proteins (MSP). The original MSP was based on human apolipoprotein A-1 (ApoA1) with the globular N-terminal domain removed.^1^

One of the primary advantages of MSP-based nanodiscs is the ability to engineer the MSP sequence to alter the properties. For example, MSP proteins have been truncated or extended to yield nanodiscs with diverse sizes from 7 nm to 17 nm.^15,16^ By linking the N- and C-termini of the MSP protein, covalently circularized nanodiscs have been developed to improve the stability and monodispersity of nanodiscs, and they enabled nanodiscs up to 100 nm.^17^ Circularized nanodiscs can be more challenging to purify and result in lower yields,^18^ which led to the development of spMSP1D1 using the SpyCatcher-SpyTag system for *in vivo* circularization.^18^ Beyond changing the size, the MSP protein has also been fused with a proximity labeling enzyme to determine binding of soluble proteins to lipid or membrane proteins inside the nanodiscs.^12^

Another useful approach to alter the properties of nanodiscs has been to design unique spectroscopic signatures. Sligar and coworkers have engineered the MSP sequence to change the tryptophan and tyrosine content.^19^ Dark MSP had all tryptophan residues mutated, and ultra-dark MSP had all tryptophan and tyrosine residues mutated. These constructs allow spectroscopy to probe proteins or peptides interacting with nanodiscs without interferences from the MSP belt. In contrast ultra-bright MSP added tryptophan residues to enhance the absorbance, which improves sensitivity for analysis techniques.

Nanodiscs have also been modified to include fluorescent reporters for binding studies. The first experiments used a D73C mutant of MSP1D1 and covalent labelling to attach a fluorophore on that single cysteine.^20^ These constructs have been used for fluorescence quenching binding assays, where labelled nanodiscs with embedded rhodopsin quench the emission of fluorescently labeled arrestin when it binds.^11^ Another application used a fluorescence resonance energy transfer (FRET) assay to measure binding of talin head domain to D73C nanodiscs.^20^ This approach is highly flexible, but the fluorescent ligands can be costly and poorly soluble. Moreover, the chemical labeling cannot be performed after assembling membrane protein nanodiscs that may contain surface exposed cysteines.

Recently, fluorescent nanodiscs have been designed based on circularized split GFP (spGFP). This approach enabled easy expression of fluorescent MSP without the need for costly ligands. spGFP nanodiscs were used with FRET to study binding of synaptotagmin-1 with anionic lipid nanodiscs.^21^ However, the circularized spGFP nanodisc approach is limited to only spGFP and could not be easily extended to other fluorescent proteins. Because not all fluorescent proteins have clear split designs, the spGFP approach may not be easy to extend to applications where multiple or different colors of fluorophores are needed.

Here, our goal was to develop fluorescent MSP with a wide range of different colors. We fused MSP1D1 or MSP1E3D1 with multiple different fluorescent proteins and a HaloTag using a short linker. We hypothesized that the addition of these C-terminal fusion proteins would not affect the formation of nanodiscs but instead to the outside of the nanodisc core. We found that these fusions expressed well and readily formed nanodiscs with similar structure and stoichiometry to conventional nanodiscs. Overall, these new designs provide a general approach for engineering nanodiscs with a wide range of spectroscopic properties, customizable fluorescent colors, and new functions that enable a range of different experiments.

## Materials and Methods

### Materials

Amberlite XAD®-2 beads were purchased from Millipore Sigma. 1,2-Dimyristoyl-*sn*-glycero-3-phosphocholine (DMPC) and 1,2-dimyristoyl-*sn*-glycerol-3-phspho-(1’-rac-glycerol) (DMPG) were purchased from Avanti Polar Lipids. One Shot BL21 Star (DE3) competent cells were purchased from Thermo Fisher Scientific. Costar Spin-X 0.22 µm filters, Halt Protease Inhibitor, and Pierce BCA Assay were purchased from Fisher Scientific. HisTrap FF 16/10, Superdex 200 Increase 10/300 GL, and Superose 6 Increase 10/300 GL columns were purchased from Cytiva. Gel filtration SEC standards were purchased from BioRad.

### Plasmid Design

MSP1D1 and MSP1E3D1 were fused with fluorescent proteins using simple linkers at the C-terminus. The rationales for the specific proteins chosen are described in the Supporting Methods. Each construct retained the 7×His tag on the N-terminus, TEV cleavage site, and MSP sequence from the original Sligar MSP plasmids.^15^ C-terminal fluorescent proteins were added with either a GS or GGGGSGGGGS linker. We chose the short GS linker to avoid potential proteolytic cleavage between the two proteins during expression, which we have observed with other proteins in *E. coli* in the past. The longer linker was included to test if additional flexibility could be added without sacrificing proteolytic stability. Full sequences and origins of each protein are provided in Table S1.

Plasmids were synthesized and cloned into the pET-28a(+) vector by GenScript and verified by sequencing. Plasmids are available to the community at Addgene (https://www.addgene.org/Michael_Marty/), and detailed information for plasmids is included in Table S1.

### Protein Expression and Purification

MSP expression and purification were carried out as previously described^1,16^ with modifications. Detailed methods are provided in the Supplemental Methods. Briefly, plasmids were transformed into BL21 DE3 cells and grown as previously described.^1,16^ MSPs were purified via immobilized metal affinity chromatography (IMAC) with standard procedures^1,16^ except in low light conditions to prevent photobleaching. After IMAC, MSPs were incubated with tobacco etch virus (TEV) protease to cleave the polyhistidine tags and were purified by SEC to separate the TEV protease from the cleaved MSP and remove any truncated fragments. After SEC, we used a BCA assay to determine the total protein concentration. Additional experimental details on SDS-PAGE characterization; absorbance and fluorescent spectroscopy; mass spectrometry; and HaloTag labeling are provided in the Supplemental Methods. Raw data are available at https://doi.org/10.5281/zenodo.18988631.

To demonstrate incorporation of an integral membrane protein, *E. coli* aquaporin Z (AqpZ) with a C-terminal sfGFP was expressed and purified as previously described.^22^ For FRET analysis, an N-terminal fusion of sfGFP to human annexin V was designed and synthesized by GenScript, with the sequence reported in Table S1. Details on its expression and purification are described in the Supporting Methods.

### Nanodisc Assembly

Nanodiscs were assembled as previously described.^1^ All samples were prepared with DMPC lipids except where noted as DMPG for FRET assays. Briefly, lipids were dissolved in chloroform, and their concentration was determined by phosphate analysis. Lipids were then dried and resolubilized in a 100 mM cholate solution to a 50 mM final lipid concentration. MSP was mixed with lipids. For the MSP1D1-based constructs, the published ratio for MSP1D1 and DMPC (1:80) was used,^1,15,16^ and MSP1E3D1-based constructs used the published ratio of 1:150.^15^ Lipids and MSP were incubated at room temperature (23 °C) on an orbital shaker for 30 mins. Then, Amberlite XAD®-2 beads were added at 50% w/v and incubated for 4 hours. Beads were removed with a 0.22 µm Costar spin filter by centrifugation at 10,000×g for 2 minutes. After filtration, nanodiscs were purified by SEC using a Cytiva Superose™ 6 increase 10/300 GL column with 200 mM ammonium acetate as the mobile phase at a flow rate of 0.5 mL/min. SEC standards were used to determine the Stokes diameter. At least three replicates were prepared for each nanodisc, which were independently analyzed by SEC, MS, and mass photometry (MP). Experimental methods for mass photometry are described in the Supplemental Methods. AqpZ nanodiscs were prepared with DMPC as previously described^22,23^ using the MSP1E3D1-sfCherry3C construct.

### Native MS and Charge Detection-MS of Nanodiscs

Nanodiscs were diluted to a final concentration of 1–3 µM in 200 mM ammonium acetate and ionized with a similar approach as for free MSPs (see Supplemental Methods) and as previously described.^24–27^ Key instrumental parameters are included for both native MS and CD-MS in Table S2 and S3. Native MS was used to determine the stoichiometry of MSP per nanodiscs by using molecular mass defect analysis^28^ in UniDec. CD-MS was used to determine the lipid to MSP stoichiometry ratio. Data was deconvolved using UniDec^29^ and UniDecCD,^30^ respectively. Additional details on MS data analysis are provided in the Supplemental Methods.

### FRET Analysis

For the FRET assay to monitor annexin V binding to nanodiscs, nanodiscs were prepared with either DMPG or DMPC using the MSP1D1-sfCherryS construct. Nanodiscs were mixed with sfGFP-annexin V to final concentrations of 0.5 µM and 1 µM, respectively. To initiate binding, calcium chloride was added to a final concentration of 8 mM. Controls were prepared without calcium. Triplicate samples were incubated in a Corning black opaque 96-well plate for 5 minutes, and measurements were done on a BioTek Synergy H1 microplate reader. The wavelengths used for FRET were 488 nm excitation and 610 nm emission. Details on FRET analysis of AqpZ nanodiscs are provided in the Supplemental Methods.

### High-speed Atomic Force Microscopy

High-speed atomic force microscopy (HS-AFM; MSNEX, RIBM, Japan) images were obtained in liquid in tapping mode operation using an AFM tip (USC-F1.2-k0.15, Nanoworld) with 0.15 N/m stiffness. In general, a 1.5 μL aliquot of each diluted sample was deposited onto freshly cleaved mica, incubated for 3 mins, and rinsed with imaging buffer prior to imaging. The imaging buffer consisted of 20 mM HEPES, 150 mM NaCl, and 5 mM CaCl_2_. The concentrations of MSP1D1 and MSP1D1-sfGFP nanodiscs used for HS-AFM studies were 30 nM and 50 nM, respectively. Particle areas for monomeric nanodiscs were quantified in ImageJ using height-based segmentation with a threshold of ∼2.2 nm to isolate individual particles. This threshold was selected to define the projected particle area while minimizing low-height peripheral signals that are most susceptible to tip-broadening effects. Center-to-center distances between interacting nanodiscs were measured in Fiji using TrackMate with a Laplacian of Gaussian (LoG) detector and an estimated particle diameter of ∼20 nm.

## Results and Discussion

### Protein Expression, Purification, and Characterization

Our goal was to develop versatile fluorescent nanodiscs, so we fused the sequence of MSP1D1 with C-terminal fluorescent protein through a simple linker. After designing the plasmids, we first evaluated protein expression. Interestingly, we noticed colonies were visibly colored during the initial transformation into BL21 *E. coli*, indicating leaky expression of the plasmid prior to induction. We experimented with growth and induction conditions, but the standard growth and induction conditions^1^ for MSP continued to give the best yields. Using lower temperature and longer induction (overnight at 18 °C) or using lower IPTG, both to slow expression, yielded more proteolytic degradation of the MSP and more free fluorescent proteins. Both control MSP1D1 and these new fluorescent MSP constructs yielded around 12 mg of purified protein per liter of culture. However, the optimal conditions for MSP expression may not give enough time for full maturation of some of the fluorescent proteins, as discussed below.^31^ Longer times may be used to improve fluorescence, but we opted to use shorter times to avoid any proteolysis of the MSP.^32^

Because our fluorescent nanodisc constructs retained the N-terminal poly-histidine tag of conventional MSPs, purification of fluoro-MSPs was similar to conventional MSPs. However, we modified the purification slightly in one aspect. Typically, MSP is purified by IMAC and is then optionally followed by TEV protease cleavage and reverse IMAC. This conventional purification scheme cannot separate truncated MSP that has lost the fluorescent protein from the full-length protein. Because the full-length protein was significantly larger than the truncated MSP, SEC was used to remove any truncated protein. Initial tests revealed that there was minimal truncated MSP in most batches, but the SEC was also useful to remove TEV and avoid overnight dialysis to exchange the protein into storage buffer. Thus, we typically performed IMAC as the first separation but replaced reverse IMAC with SEC for purification after TEV protease cleavage. One limitation of this approach is that it will not separate missed cleavage that retained the His-tag, but we did not observe any significant missed cleavage in our MS data (Figure S1).

SDS-PAGE analysis (Figure 1) revealed similar purity in fluoro-MSPs to conventional MSP. General protein staining showed a single dominant band for our controls MSP (Figure 1A). Fluoro-MSPs also had mostly clean bands that appeared slightly lower than the expected molecular weight. For example, MSP1D1-sfGFP has a mass of 48 kDa (Table S4) but appears with a primary band around 37 kDa and a minor band around 48 kDa. Because the mass measured by mass spectrometry (see below) agrees with the predicted mass, our interpretation is that the covalently crosslinked fluorophore core in the fluorescent protein resists denaturation and thus has a more compact shape that causes faster migration in the gel relative to standards. We hypothesize that the minor band closer to the expected molecular weight is from a subset of proteins where the fluorophore failed to mature and thus denature more in SDS. Supporting this hypothesis, the HaloTag construct lacks both the upper band and faster travel. Moreover, only the primary bands appear on fluorescent imaging (Figure 1B).

**Figure 1.**
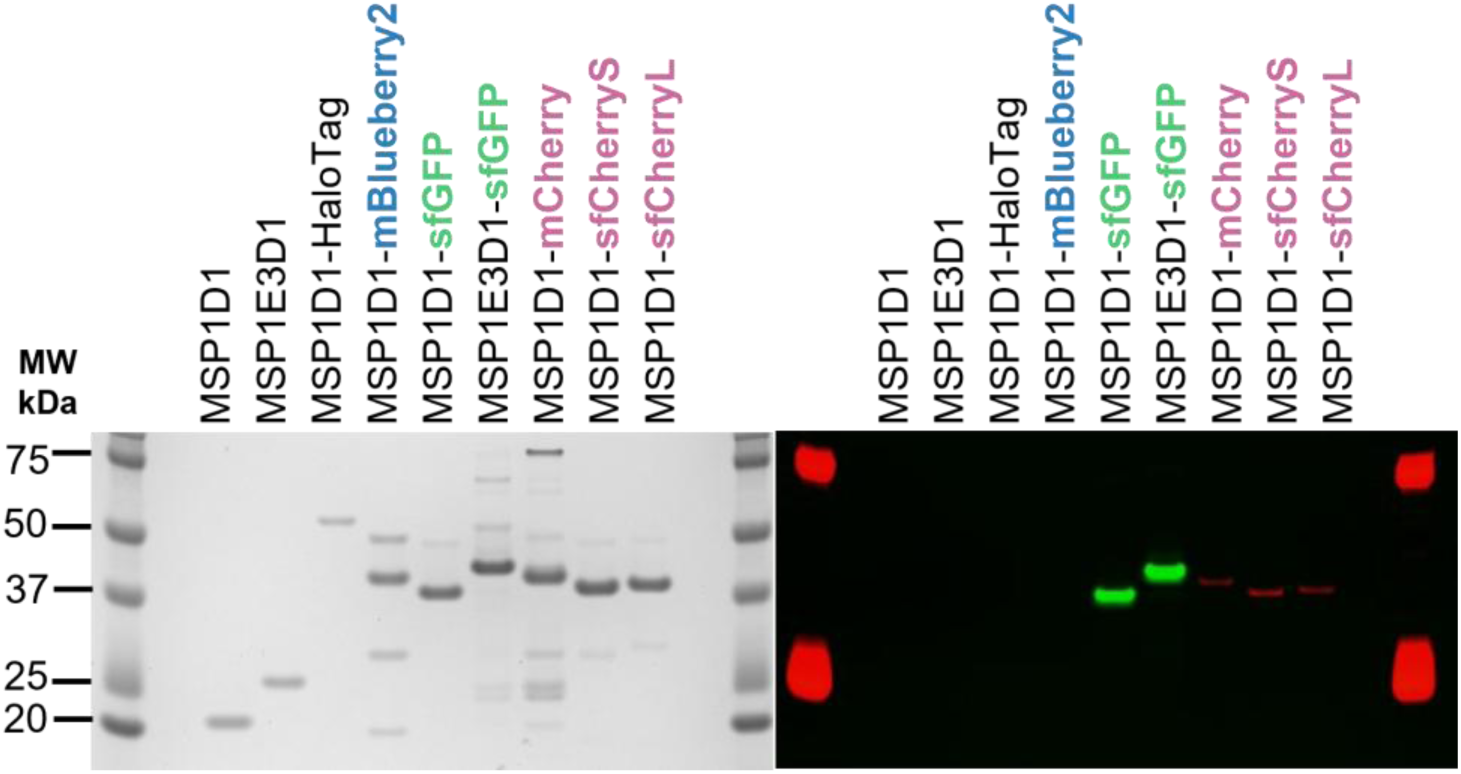
SDS-PAGE of MSPs with either Imperial™ Protein Stain (*left*) and with background subtracted fluorescent imaging (*right*) in either red (Ex 550/Em 602/50) or green (Ex 488/Em 528/32). Unfortunately, the blue fluorescence filters available were not low enough to excite mBlueberry2, but fluorescence is observed in solution. The construct names are included above each lane, and standards are in the first and last lane.

Interestingly, the superfolder proteins show less of the immature upper band, indicating more efficient formation of the fluorophore. Moreover, the non-superfolder proteins, with mBlueberry2 and mCherry, had lower molecular weight bands, which could indicate truncation. Our interpretation is that the superfolder constructs are less susceptible to proteolytic cleavage during expression and purification because they more quickly fold into a stable structure. We also observed that sfGFP constructs show more intense fluorescent bands than the mCherry-based constructs, which was expected due to the lower efficiency and quantum yield of mCherry-derived fluorescent proteins.^33^

To confirm the integrity of the protein sequence, we performed intact mass analysis on the purified proteins (Figure S1). All fluoro-MSPs had mass values that were close to the sequence mass but lower by 17–23 Da (Table S4). This mass difference reflects the loss of two protons (-2 Da) and water (-18 Da) during chromophore formation.^34^ For example, the chromophore formation reaction for sfGFP has a reported mass loss of 20 Da,^35^ which aligns with our measured mass. Only small peaks were observed for truncated products, with the highest being around 6% in the mCherry construct. These results confirm the expected protein mass and formation of the fluorophore. Finally, we also characterized the spectral properties of our fluorescent MSP by measuring their respective fluorescent excitation and emission (Figure S2A) and their UV-visible absorption (Figure S2B).

### Optimizing and Validating Nanodisc Assembly

After successful expression and purification of fluoro-MSP proteins, we assembled nanodiscs using these new constructs. Fluorescence spectra of nanodiscs (Figure S3) closely matched those of the MSP. The primary challenge in working with fluoro-MSPs was accurately determining their concentration. We could not rely solely on the absorption at 280 nm to determine the protein concentration because each chromophore absorbed inconsistently at 280 nm due to photobleaching and incomplete maturation. Thus, we used a BCA assay to estimate the total protein concentration. In general, using the BCA concentration directly yielded nanodiscs with the proper loading. However, it may be necessary to empirically optimize the assembly ratio, and we recommend testing it on the first use of each batch.

We first examined the SEC data during nanodisc purification (Figure 2). As expected, fluoro-MSP nanodiscs eluted earlier than controls because the added fluorescent proteins increased the hydrodynamic radius of the complex. From the peak retention volume, we calculated the Stokes diameter of the nanodiscs (Figure S4). Control nanodisc had a Stokes diameter of 9.3 ± 0.2 nm, which agreed with literature values.^1,15^ The MSP1D1-sfGFP nanodisc had a Stokes diameter of 13.6 ± 0.1 nm (Table S5), which was similar to the previously-described spGFP circularized nanodiscs.^21^ Other fluorescent nanodiscs had similar diameters in the 11–14 nm range (Table S5). The peak shapes were generally Gaussian, indicating a relatively monodisperse size distribution. Overall, these data indicate that fluorescent fusions extend the size of the nanodisc and support a model where the fluorescent protein is on the side of the core disc.

**Figure 2.**
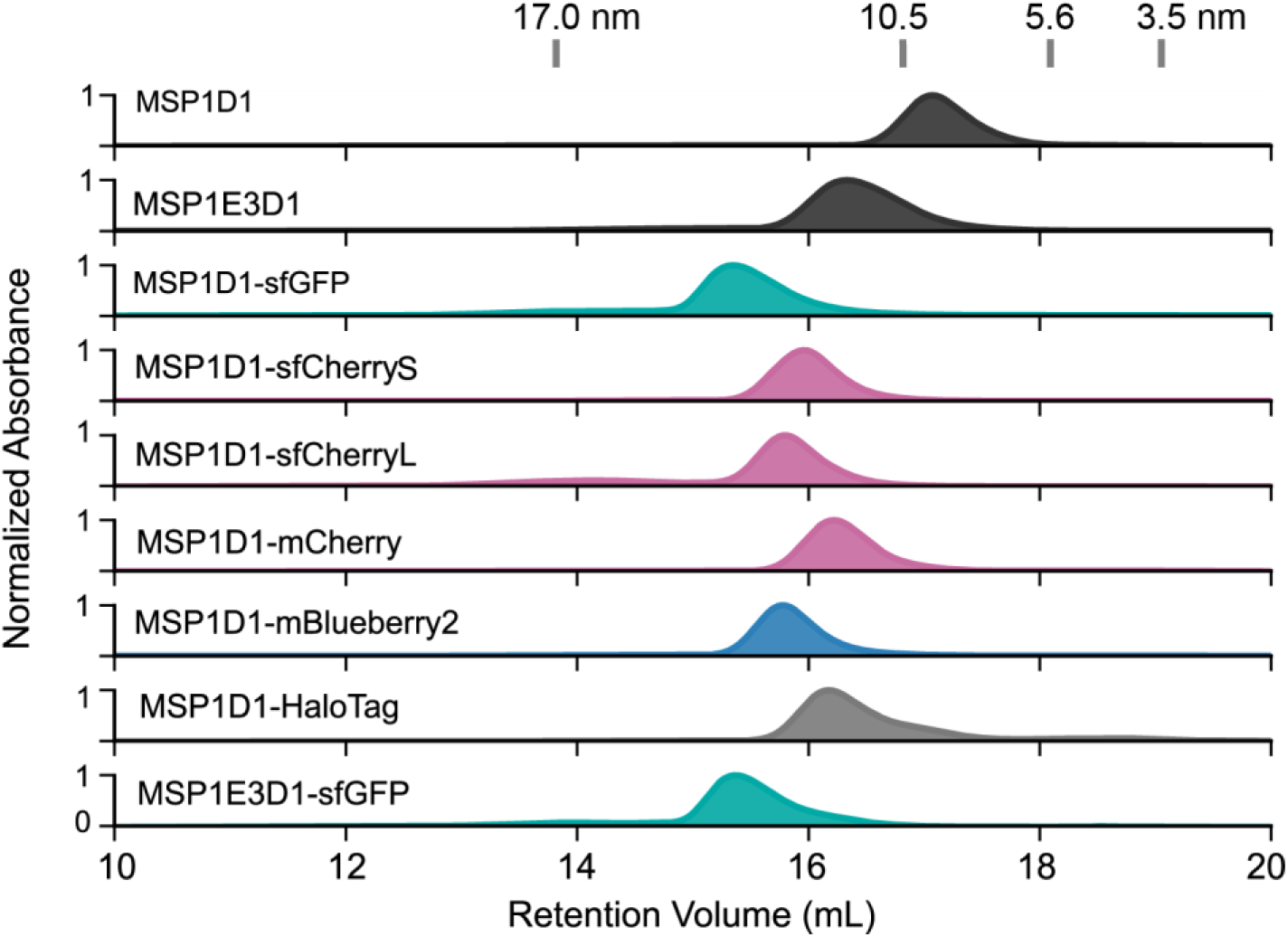
Size exclusion chromatograms of nanodiscs made (listed from *top* to *bottom*) from control MSP1D1 (*black*), MSP1E3D1 (*black*), MSP1D1-sfGFP (*green*), MSP1D1-sfCherryS and MSP1D1-sfCherryL (*red*), MSP1D1-mBlueberry2 (*blue*), MSP1D1-HaloTag (*grey*), and MSP1E3D1-sfGFP (*green*). SEC standards are marked at the top with their Stokes diameter annotated at their respective retention volumes. Each nanodisc was assembled in triplicate, and a single representative chromatogram is shown for each. Note, these data are from the primary purification of nanodiscs following assembly and are not reinjected for analysis.

Interestingly, despite having a significantly larger mass and the expected number of lipids (see below), MSP1E3D1-sfGFP had a similar measured Stokes diameter to MSP1D1-sfGFP. We propose that some of these observed variations in Stokes diameter could arise from register shifts of the two MSP belts in the nanodisc, which could position the two fluorescent proteins in different relative angles depending on the specific fusion protein and MSP. Additional evidence for hypothesis is presented below from HS-AFM, and a more detailed discussion is provided in the Supplemental Discussion.

After SEC revealed that the nanodiscs were monodisperse and approximately the right size, we quantified the stoichiometry of the nanodiscs with mass spectrometry. Using native MS and mass defect analysis,^22,23,28^ we measured the masses of the mostly intact nanodiscs and confirmed that nanodiscs had only two MSP belts per complex (Figure S5). However, native MS requires some collisional activation to resolve nanodiscs, which risks dissociating some lipids and underestimating the lipids per complex.

To more accurately measure the number of lipids per complex, we used CD-MS, which measures the charge and *m/z* of each ion simultaneously to get a mass distribution without needing to add collisional energy to resolve the nanodisc (Figure S6 and S7).^36^ With CD-MS, we observe that the fluorescent nanodiscs are consistently larger, but the mass shifts can be explained by the additional mass of the fluoro-MSP proteins. When we subtract the mass of the two belt proteins, we determined that fluorescent nanodiscs from the MSP1D1 constructs had a similar lipid fill ratio to that of the control MSP1D1 nanodiscs (Figure 3). The other fluoro-MSPs followed a similar trend, where the stoichiometry matched conventional MSPs (Figure S6). To validate the CD-MS results, we performed mass photometry, which also showed the expected mass shifts for addition of fluorescent proteins and consistent nanodisc filling (Figure S8 and Table S6). Additional discussions of the CD-MS and mass photometry results are provided in the Supplemental Discussion.

**Figure 3.**
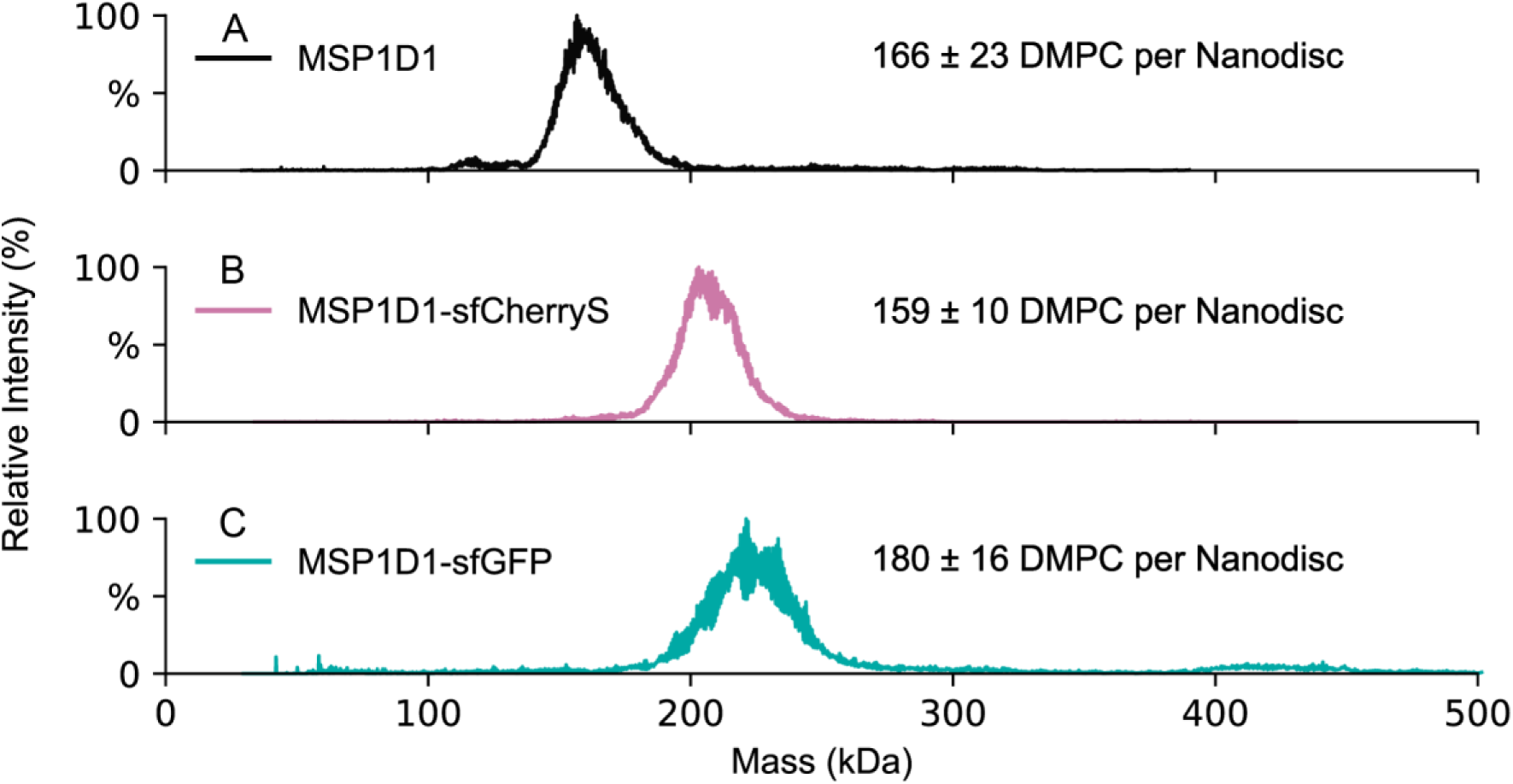
CD-MS distribution of (A) control nanodiscs (*black*), (B) MSP1D1-sfCherryS nanodiscs (*red*), and (C) MSP1D1-sfGFP nanodiscs (*green*). Data is shown for a representative replicate. The legends indicate the average and standard deviation number of DMPC lipids per nanodisc complex, which is calculated for three independent nanodisc assemblies by subtracting the mass of two MSPs and dividing by the lipid mass.

### Imaging Fluorescent Nanodiscs

We performed high-speed atomic force microscopy (HS-AFM) to image nanodiscs made from conventional MSP1D1 and from MSP1D1-sfGFP. Nanodiscs were deposited on freshly cleaved mica under buffering conditions (Figure 4). Representative HS-AFM images show distinct nanodisc particles measured in both samples. Some higher-order assemblies are observed in both samples, discussed more below (Figure 4A).

**Figure 4.**
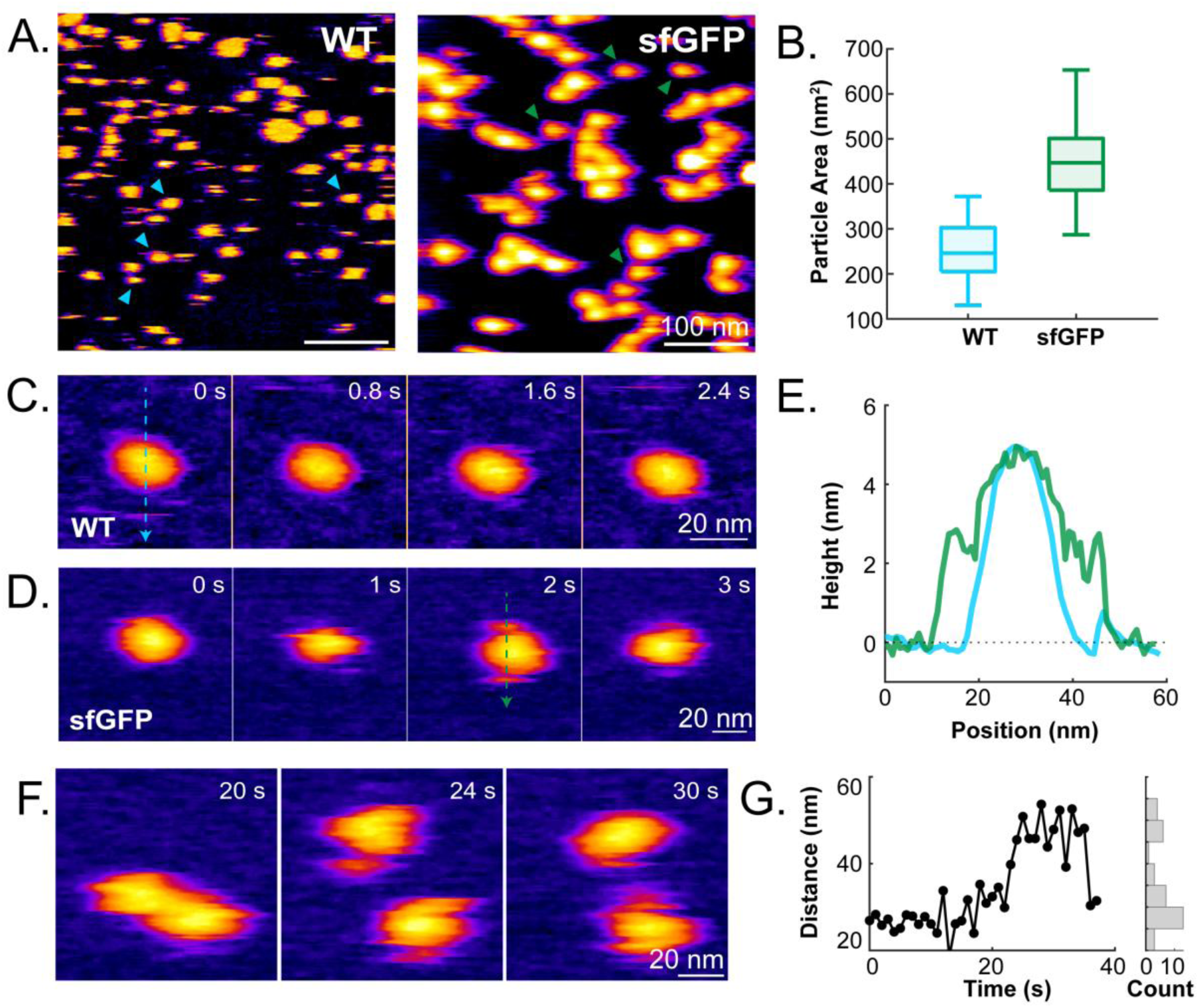
Structural characterization of nanodiscs. (A) Representative HS-AFM images of the MSP1D1 (WT) and MSP1D1-sfGFP (sfGFP) nanodiscs deposited on mica substrate. Colored arrows (*cyan* and *green*) indicate isolated nanodiscs. (B) The box-whisker plot of monomeric particle areas measured in (A) using height-based segmentation with a threshold of ∼2.2 nm to define projected particle areas for WT 252.1 ± 9 nm^2^ (*n*=20) and sfGFP 460 ± 99 nm^2^ (*n*=20), p<0.001. (C-D) Realtime structural dynamics of individual (C) WT and (D) sfGFP nanodisc. (E) The cross-sectional profiles of the WT and sfGFP nanodisc along the (C) *cyan* and (D) *green* arrow, respectively. The two fused-GFP molecules sometimes exhibit two globular shoulders at the opposite side of this nanodisc. (F) Realtime association and dissociation of two sfGFP nanodiscs observed by HS-AFM. (G) The intermolecular distance with a histogram of two sfGFP nanodiscs (center-to-center) observed in (F) over time.

Monomeric nanodiscs were further selected for quantitative image analysis, revealing that conventional MSP1D1 nanodiscs had a mean projected particle area of 252.1 ± 9 nm^2^ (*n*=20), whereas MSP1D1-sfGFP nanodiscs had a significantly larger mean area of 460 ± 12 nm^2^ (*n*=20) (Figure 4B). Assuming nanodiscs are circular geometry, these values correspond to average diameters of 17.7 ± 2.6 nm for MSP1D1 nanodiscs and 24.1 ± 2.6 nm for MSP1D1-sfGFP nanodiscs. These results are systematically higher than the Stokes diameters observed above. The larger apparent diameter of the control MSP1D1 nanodiscs may reflect partial spreading or flattening of soft discoidal lipid– protein particles on mica during HS-AFM imaging, with a minor contribution from tip convolution.

We further monitored particle morphology and dynamic behavior of individual MSP1D1 and MSP1D1-sfGFP nanodiscs (Figure 4C and 4D, Movie S1 and S2). Both nanodiscs had a round, disc-like geometry with a central bilayer thickness of 4.6 ± 0.3 nm (Figure 4E), in agreement with the expected height of a DMPC bilayer in nanodiscs.^37,38^ Notably, MSP1D1-sfGFP nanodiscs sometimes displayed two small, lobe-like protrusions (∼2.5 nm in height) positioned on opposite sides of the disc. These protrusions are consistent with the size of the two sfGFP tags^39^ fused to the MSP1D1 proteins that assemble into each nanodisc with a dimeric belt architecture.

In contrast, MSP1D1-mCherry nanodiscs had lobes positioned on the same side, around a 65° angle between each other (Figure S9). These data support a model proposed above where mCherry nanodiscs are more compact than sfGFP nanodisc due to different registers of the MSP. In both cases, these additions do not seem to cause defects in the central nanodisc core but instead move dynamically around the core.

Negative stain TEM was also used to characterize the particle sizes. Here, the average diameters were 10.4 ± 0.4 and 16.8 ± 1.9 nm, respectively (Figure S10). Unlike AFM, TEM lacked the resolution to distinguish the GFP fusions. It also has potential biases in dehydration artifacts and limited ability to detect flexible domains. Although the absolute values differ and measure different underlying properties for these dynamic and extended particles, HS-AFM, TEM, and SEC consistently show that sfGFP fusion increases the apparent nanodisc size, supporting successful formation of larger fluorescent nanodiscs.^1,40^ The exact structure and sizes are difficult to define, but these data are consistent with a model of a core nanodisc with flexible fluorescent proteins appended to the side.

For MSP1D1-sfGFP nanodiscs, the presence of sfGFP tags introduces stronger intermolecular interactions than those observed for MSP1D1. Reversible association-dissociation events between MSP1D1-sfGFP nanodiscs were frequently observed over time (Figure 4F and 4G, Movie S3). To quantify the dimeric interactions, we measured the center-to-center distance between two nanodiscs over time and constructed an accumulated distance histogram (Figure 4G). Our results show that the stable dimeric assembly had a separation distance of ∼27 ± 3 nm. Around 24 s, the two MSP1D1-sfGFP nanodiscs separate into distinct monomers, each showing two visible sfGFP features, and subsequently reassociate through sfGFP-driven interactions, reforming the dimer-like arrangement. The reversibility of these interactions further supports a model of fluorescent proteins on the outside of the nanodisc rather than the GFP fusing with the membrane, which would likely be irreversible.

Although fluorescent nanodiscs are more likely to dimerize on the imaging surface, we do not observe significantly higher amounts of dimer at more dilute concentrations typically used for solution assays, as evidenced in CD-MS (Figure S7) and mass photometry (Figure S8). The concentration for mass photometry was similar to that of AFM loading, and we saw no significant fraction of dimer in those measurements. Thus, we interpret these as effects only observed in surface-absorbed conditions and not in bulk solution.

### Application to FRET

To demonstrate an experimental application of our fluorescent nanodiscs, we tested our new fluorescent nanodiscs in a FRET assay (Figure 5). Here, we created a construct of annexin V conjugated with sfGFP. Annexin V binds to anionic lipids in the presence of calcium and is used as a marker for apoptosis.^41,42^ Annexin is a useful system to test binding of peripheral membrane proteins to nanodiscs.^43,44^ We created nanodiscs with MSP1D1-sfCherry either DMPC or DMPG lipids. Bao and coworkers used a similar assay to with spGFP nanodiscs binding an mCherry tagged protein,^21^ the inverse of our approach.

**Figure 5.**
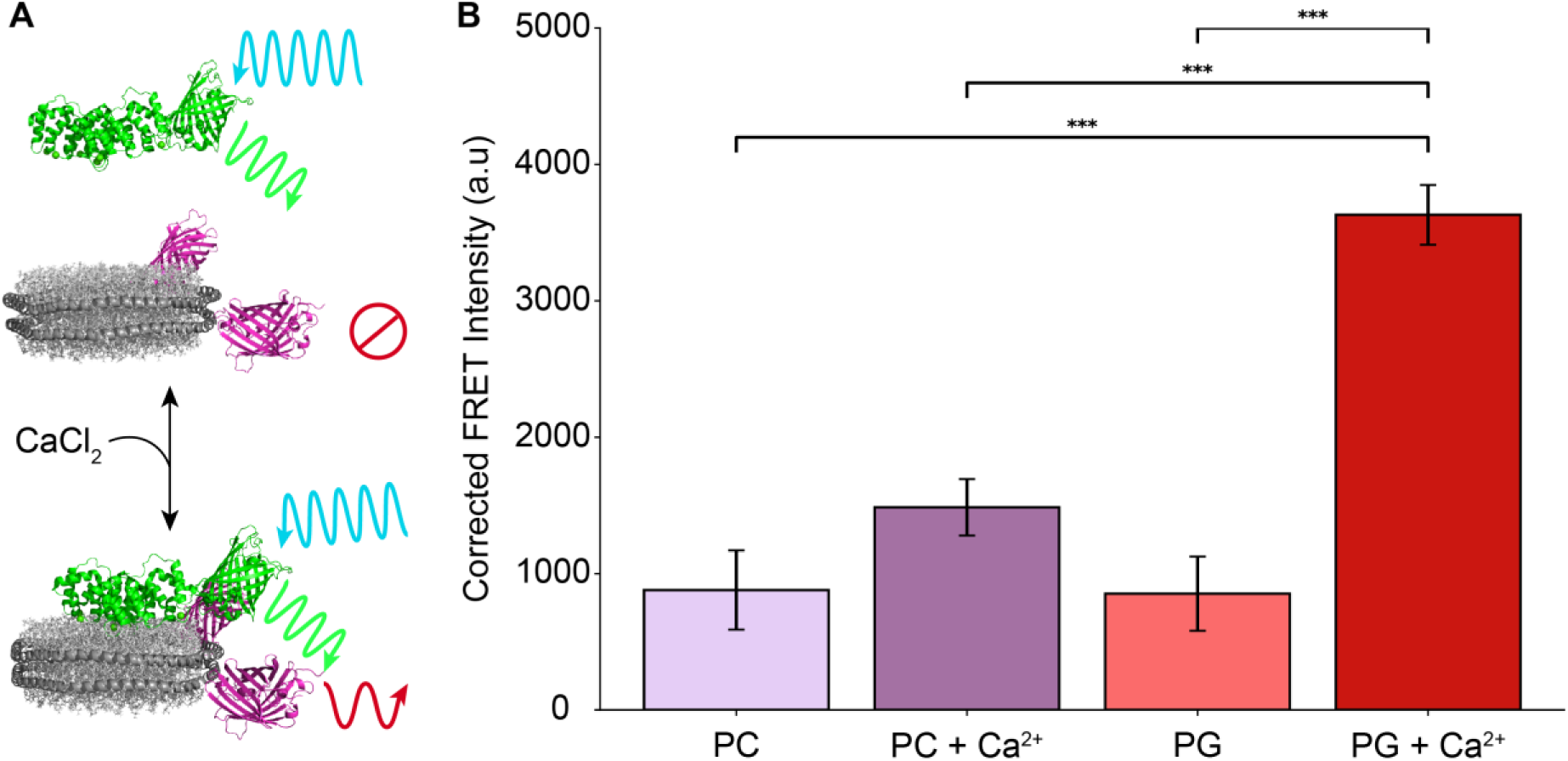
(A) Schematic of sfGFP-annexin V binding to sfCherry nanodiscs in the presence of anionic lipids and calcium. Models are only approximations to illustrate the concept. (B) Corrected FRET intensity at excitation 488 nm and emission 610 nm for annexin V added to nanodiscs with either DMPC or DMPG lipids and with or without calcium. Statistical significance was calculated from replicate nanodisc assemblies (*n*=3) using a standard one-way ANOVA followed by a Tukey’s HSD post hoc analysis. (*** p<0.001).

Monitoring the FRET intensity, lower FRET intensities were observed for both DMPC and DMPG nanodiscs in the absence of calcium. Upon addition of calcium, significantly stronger FRET intensity was observed for anionic DMPG nanodiscs but not for zwitterionic DMPC nanodiscs (Figure 5B), demonstrating specific membrane interactions. These data demonstrate the use of our fluorescent nanodisc designs to monitor membrane binding for a diverse color pallet.

FRET can also be used to monitor membrane protein incorporation into nanodiscs. Here, we assembled nanodiscs with AqpZ tagged with sfGFP into nanodiscs with sfCherry-tagged MSP. FRET was observed in nanodiscs containing the AqpZ but not in controls that lacked AqpZ (Figure S11). SDS-PAGE and MP also confirmed incorporation. Together, these data demonstrate that fluorescent MSP fusions can be used for integral membrane protein nanodiscs, and the ability to have distinct colors on the MSP is useful in tracking both proteins and confirming incorporation. A more extended discussion is provided in the Supplemental Discussions.

### Design Considerations

In the prior sections, we demonstrated that fluorescent protein fusions form effective nanodiscs and can be applied in different ways. Here, we will discuss potential considerations in design of fluoro-MSP sequences. The designs discussed above all had a simple glycine-serine linker that worked well. We chose this initial design because we wanted to limit the possibility of proteolytic cleavage between the MSP and fusion protein. However, we also tested a longer linker, GGGGSGGGGS, to see if it was more susceptible to cleavage, and it had no noticeable effect on expression or nanodisc assembly (Figures 1 and S4). This initial observation suggests that the linker may not be a critical feature of the design, but additional studies will be necessary to explore whether other types of linkers and longer lengths are feasible.

Another design consideration is whether “superfolder” versions of the fluorescent proteins had any benefit over standard fluorescent proteins.^45^ Here, we tested an mCherry fusion against the sfCherry version. As with the linkers, there was no noticeable difference between the two in nanodisc assembly. However, Figure 1 reveals that the superfolder versions yielded somewhat purer protein with less immature fluorophore. We experimented with longer maturation times as described above and encountered some issues with protein degradation, which likely limits the use of slower forming fluorophores.^31^ Thus, we recommend faster folding fluorescent proteins like sfGFP and sfCherry to prevent protein degradation.^46^

Finally, we considered how flexible the design was for different protein constructs. We successfully expressed and formed nanodiscs with a range of fluorescent proteins at different colors, including GFP, mCherry, and mBlueberry2. We also created a HaloTag7 fusion, which is an enzyme that covalently attaches halogenated compounds to itself. Halo-MSP was easier to work with than fluorescent MSPs because its concentration can be measured directly by 280 nm absorbance, like conventional MSPs. We confirmed that the HaloTag was still catalytically active in both the free MSP form and in the nanodisc. Both achieved around 70% labeling with a HaloTag-coumarin ligand (Figure S12). The highly selective labeling of nanodiscs via HaloTag opens new opportunities to selectively label samples after the nanodiscs are formed without perturbing membrane proteins inside. Overall, these data demonstrate that diverse fusions can be created that give a range of new functionality to nanodiscs. In particular, the Halo-MSP construct enables a diverse range functional handles to be attached to nanodiscs.

### Conclusions

Here, we have engineered diverse MSPs fused with fluorescent proteins and a HaloTag. We demonstrated that these new MSP form nanodiscs with two fluorescent proteins per nanodiscs while maintaining similar stoichiometries and structures as conventional nanodiscs. We have also demonstrated that these fluorescent MSPs are suitable for assembly of membrane protein nanodiscs. Our primary goal in this manuscript was to evaluate the feasibility of nanodisc design with these different constructs and to provide these plasmids as an open resource for the community.

To demonstrate a potential application of these fluoro-MSPs, we have performed FRET to detect peripheral membrane protein binding to lipids in the nanodiscs. However, we imagine a number of potential future applications, from single molecule fluorescence experiments to FRET. Outside of their utility in direct analytical assays, our lab has also found that fluoro-MSPs are very useful in monitoring assembly of membrane protein nanodiscs, as shown in the example presented here with assembly of AqpZ. By using the sfCherry MSP with a GFP tag on the target membrane protein, both the MSP and target membrane protein can be independently monitored visually and spectroscopically to track nanodisc assembly and purification. FRET can also be used to confirm co-incorporation.

The Halo-MSP presents especially wide opportunities to functionalize nanodiscs for future studies, such as with biotinylation,^47^ proximity labeling,^48^ or specialized ligands for targeting. Due to its biorthogonal chemistry, a single batch of nanodiscs could be split and functionalized separately after nanodisc assembly without modifying embedded membrane proteins. Our results indicate a wide application space to explore in covalent modification of nanodiscs with unique functional properties.

## Supporting Information

Additional detailed information included in the supporting information: Supplemental Methods for plasmid design, MSP expression and purification, SDS-PAGE, spectroscopy, MS, HaloTag labeling, expression and purification of annexin V, MP, and FRET of AqpZ nanodiscs. Captions for Supplemental Movies. Extended discussions of results and membrane protein incorporation. Supplemental Figures of MS results, fluorescence and absorbance spectra, Stokes diameters, AFM and TEM images, MP results, AqpZ nanodisc characterization, and Halo-MSP labeling. Supplemental Tables for each fused MSP sequence, relevant mass spectrometer settings, measured mass data, and Stokes diameters.

## Acknowledgements

The pMSP1D1 and pMSPE3D1 plasmid was a gift from Stephen Sligar (Addgene plasmid #20061 and #20066). The fluorescent protein sequences were provided by FPbase. The HaloTag sequence was adapted from the Los *et al.* sequence (https://doi.org/10.1021/cb800025k). The authors thank Oluwaseun Fapohunda, who designed plasmids and performed preliminary work that demonstrated feasibility. The authors thank the Samanta Lab and Que Lab at the University of Texas at Austin for allowing use of their gel imaging system and fluorimeter, respectively. TEM imaging was performed by Michelle Mikesh at the Center for Biomedical Research Support Sauer Structural Biology Lab at the University of Texas at Austin. This work was funded by National Institutes of Health and National Institute of General Medical Sciences (R35 GM128624 to MTM and R35 GM150528 to YCL) and Beckman Scholar Program to IK. Raw data from SEC, spectroscopy, TEM, SDS-PAGE, MP, and MS are available at https://doi.org/10.5281/zenodo.18988631.

## Author Contributions

FAM and ARW designed plasmids. IK performed preliminary experiments. EC, ARW, and SC expressed and purified proteins, assembled nanodiscs, and performed analyses. BHT and EC developed the MS methods. High-speed AFM data was performed by SBL and NPE, supervised by YCL. The manuscript was written by EC, ARW, and MTM with edits from other co-authors. The project was supervised by MTM.

## Notes

The authors declare no conflict of interests.

