## Supporting Information for "Design of Fluorescent Membrane Scaffold Proteins for Nanodiscs"

### Supplemental Methods

#### Plasmid Design

We chose these fusion proteins for several reasons, and our design evolved in response to our ongoing experimental observations. First, we included the superfolder green fluorescent protein (sfGFP) because it is highly stable and commonly used in our lab and in others.[1] Next, we sought to expand the color palette and chose mCherry[2] and mBlueberry2[3] as two common fluorescent proteins with different colors. However, after observing initial problems with the stability of both constructs, we chose to use a superfolder version of mCherry, sfCherry.[4] In later constructs designed for MSP1E3D1, we further changed the sfCherry to the sfCherry3C form to improve brightness.[5] Finally, to enable a much wider range of fluorescent probes and other handles to be added, we designed a construct with a HaloTag fusion. Here, the Halo-MSP offers the possibility of adding different chemical tags and doing highly specific labeling at any stage of protein expression and nanodisc assembly.

#### MSP Expression

For protein expression, we followed previously described methods.[6] Briefly, plasmids were transformed into One Shot™ BL21 Star™ (DE3) cells. Singular colonies were selected and grown overnight in 100 mL Fisher Bioreagents™ LB Broth supplemented with 0.1 mg/mL kanamycin. These starter cultures were inoculated into 1 L flasks of Fisher Bioreagents™ Terrific Broth in Thomson Instrument Company Ultra Yield flasks, and cells were grown at 37 °C and 200 rpm until the optical density at 600 nm reached 0.6–0.8. Cultures were induced with 1 mM IPTG and grown for an additional 3 hrs at 37 °C. Cells were harvested and frozen at −80 °C.

All cell lysis was performed in low light conditions to mitigate photobleaching. Cells were resuspended in lysis buffer (20 mM Tris, 150 mM NaCl, 5 mM β-mercaptoethanol (BME), 1% Triton X-100, 1:100 Halt Protease Inhibitor, pH 7.4) and lysed by either homogenization on a Microfluidics (LM20) microfluidizer at 20,000 psi for 5 cycles or by sonication at 40% amplitude with a 3-minute cycle at 30 s on-off intervals with modified lysis buffer (20 mM Tris, 150 mM NaCl, 5 mM BME, 1% Triton X-100, 1:100 Halt Protease Inhibitor, 1 mg/mL lysozyme, 10 µg/mL DNase, pH 7.4). Lysate was then clarified at 20,000 rpm for 30 mins.

#### MSP Purification

Protein purification was similar to published methods[6,7] but done in low light conditions to avoid photobleaching. After clarification, lysate was passed through a 0.45 µm filter, and filtered lysate was loaded onto a Cytiva HisTrap FF 16/10 column equilibrated in loading buffer (40 mM Tris, 300 mM NaCl, 1% Triton X-100, 5 mM BME, pH 8.0). The column was then washed in the loading buffer for at least two column volumes (CV). Then, it washed with two CV of the first wash buffer (40 mM Tris, 300 mM NaCl, 50 mM sodium cholate, 20 mM imidazole, and 5 mM BME, pH 8.0) followed by two CV of the second wash buffer (40 mM Tris, 300 mM NaCl, 50 mM imidazole, and 5 mM BME, pH 8.0). The protein was then eluted with elution buffer (40 mM Tris, 300 mM NaCl, 400 mM imidazole, and 5 mM BME, pH 7.4).

Eluent was then incubated with tobacco etch virus (TEV) protease at a 1:50 mass ratio overnight. After His-tag removal, each protein was concentrated on a 30 kDa MWCO filter and loaded onto a Cytiva Superdex 200 Increase 10/300 GL for size exclusion chromatography (SEC) in a buffer of 20 mM Tris, 150 mM NaCl, and 5 mM BME, pH 7.4. Fractions from SEC were analyzed by SDS-PAGE to verify the correct molecular weight. Finally, we used a Pierce™ BCA Assay (Thermo Fisher Scientific) to determine total protein concentration.

#### SDS-PAGE

Samples for SDS-PAGE were treated with dithiothreitol and diluted into Laemelli sample buffer. They were not boiled to keep the fluorescent protein intact for in-gel fluorescence. SDS-PAGE was performed at 120 V for 60 mins using a 12% Tris gel. In-gel fluorescence was visualized with BioRad ChemiDoc Imaging system. Then, the gel was stained with Imperial™ Protein Stain (Thermo Fisher Scientific) for 15 mins on a rocker. Destaining was carried out with deionized water overnight and imaged with BioRad ChemiDoc Imaging system.

#### Absorbance and Fluorescence Spectroscopy

For absorbance spectroscopy, purified MSP protein samples were diluted to 5 μM based on the concentration measured by BCA analysis. Then, each sample was measured in triplicated on a BMG Labtech Spectrostar Nano using an Eppendorf UVette with a path length of 1 cm. Spectra were measured from 220–1000 nm.

For free MSP and nanodisc fluorescence characterization, all samples were diluted to 1 μM, and the fluorescent excitation and emission were measured on an Agilent Technologies Cary Eclipse Fluorescence Spectrophotometer. Fluorescence excitation and emission were measured in cuvettes at each fluorescent protein's respective excitation and emission wavelengths.[8]

#### Mass Spectrometry of MSP Proteins

MSPs were further characterized with mass spectrometry (MS) following previously described methods.[9] Briefly, samples were buffer exchanged into 200 mM ammonium acetate with Micro Bio-Spin P-6 columns (BioRad). Then, nano-electrospray ionization was performed in positive ionization mode using borosilicate tips pulled using a P-1000 micropipette puller (Sutter Instruments). Samples were analyzed on a Thermo Q Exactive HF UHMR, and key instrument settings are reported in Table S2. Data was deconvolved with UniDec as described below.[10]

#### Mass Spectrometry Data Analysis

Mass spectra for free MSP constructs were deconvolved in UniDec using default settings with a charge range of 1 to 25 and a mass range of 1,000 Da to 60,000 Da. Native mass spectra for nanodiscs were deconvolved with the Nanodisc DMPC preset except with a FWHM setting of 4 Th. CD-MS deconvolution was performed in UniDecCD with the default settings except for the charge range was set from 1 to 50, *m/z* spread FWHM of 10, and charge spread FWHM of 4.

#### HaloTag Labeling

Halo-MSP and Halo-MSP nanodiscs were labeled using previously described methods.[11] Briefly, HaloTag® Coumarin Ligand was purchased from Promega. A working stock was made at 100 µM in dimethyl sulfoxide (DMSO). Then, 10 µM of either Halo-MSP or Halo-MSP nanodiscs were mixed with HaloTag® Coumarin at a 1:1 mol ratio (a final DMSO concentration of 1%) and incubated for 15 mins at room temp (23 °C) with shaking set to 40 rpm. After incubation, samples were buffer exchanged into 200 mM ammonium acetate with Micro Bio-Spin P-6 columns, and samples were analyzed with mass spectrometry using methods that matched the free MSP or intact nanodiscs, as applicable.

#### Expression and Purification of sfGFP-Annexin V

Annexin V was fused with an N-terminal sfGFP and 7×His tag based on similar construct designs.[12,13] We also included a TEV cleavage site in between sfGFP and Annexin V. Plasmids were synthesized and cloned into pET-28a(+) vector by Genscript and verified by sequencing. This plasmid is also available to the community at Addgene (https://www.addgene.org/Michael_Marty/), and detailed information is included in Table S1.

For protein expression we followed similar procedures as described above with some modifications. Briefly, after the starter culture was inoculated into 1 L flasks of Fisher Bioreagents™ LB in Thomson Instrument Company Ultra Yield flasks, cells were grown at 37 °C and 200 rpm until the optical density at 600 nm reached 0.6–0.8. Cultures were induced with 1 mM IPTG and grown overnight at 18 °C.

For purification, we used similar procedures to those described above. Cells were lysed with the microfluidizer and clarified. IMAC was performed using similar buffers but without any detergents. After IMAC, SEC was performed, as described above. Peak fractions were collected and analyzed by SDS-PAGE as described above to check for in-gel fluorescence and correct molecular weight. sfGFP-Annexin V was then concentrated on a 50 kDa molecular weight cut off filter. Finally, we used MS to verify the intact mass of the sfGFP-Annexin V, as described above.

#### Mass Photometry

Mass photometry was conducted on a Refeyn Two MP, using the regular field of view (FOV), 4 × 11 µm, at 500 Hz and 60 s of acquisition. Bovine serum albumin was used for a three-point mass calibration, at monomer, dimer, and trimer values. Samples were diluted to a final concentration of 20 nM in the sample well using 200 mM ammonium acetate. Membrane protein nanodiscs were analyzed using the large FOV, 12 × 17 µm, at 135 Hz and 60 s acquisitions. Here, GroEL was added to the calibration to extend the mass range, and samples were diluted as described above.

#### FRET Analysis of AqpZ Nanodiscs

AqpZ nanodiscs were characterized with FRET analysis to demonstrate co-incorporation of sfGFP-tagged AqpZ into nanodiscs formed with MSP1E3D1-sfCherry3C. After assembly, AqpZ nanodiscs were purified from empty nanodiscs using IMAC. Both the flowthrough (containing empty) and elution (containing AqpZ) fractions were collected and purified by SEC. Each was diluted to 0.5 µM and measured by FRET as described for the annexin V.

### Captions for the Supplemental Videos

**Supplementary Movie S1.** Structural dynamics of an MSP1D1 nanodisc (Figure 4C). Imaging parameters: 0.8 s/frame, 0.25 nm/pixel.

**Supplementary Movie S2.** Structural dynamics of an MSP1D1-sfGFP nanodisc (Figure 4D). Imaging parameters: 1 s/frame, 0.25 nm/pixel.

**Supplementary Movie S3.** Association and dissociation dynamics between two MSP1D1-sfGFP nanodiscs (Figure 4F). Imaging parameters: 1 s/frame, 0.25 nm/pixel.

### Supplemental Discussion

#### Extended Discussion of SEC Results

As noted in the main text, there were some shifts in the Stokes diameter measured by SEC that appeared larger than expected. CD-MS and MP measurements confirm that the masses of the particles are very similar and close to expected values (discussed more below). Thus, these deviations either reflect different shapes or measurement artifacts.

One potential factor that could cause different Stokes diameters is register shifts in the MSP belt orientation. Crosslinking MS data has found that the primary orientation of the two ApoA-I proteins in reconstituted lipid nanoparticles in an antiparallel arrangement with helix 5 to helix 5 (5/5) overlap, which would position the C-termini close together[14] and favor a more compact structure. However, the authors note additional crosslinks that are unexplainable with the primary orientation and suggest a secondary helix 5 to helix 2 orientation, which would position the C-termini further apart in a more extended structure.

It could be that C-terminal fusion proteins bias the preferred register and shift the C-terminal to C-terminal angle differently in the MSP1D1 nanodiscs than in the MSP1E3D1 version and differently between fusion proteins. In HS-AFM data (Figure 4), sfGFP nanodiscs seem to adopt a more extended structure with GFP-to-GFP angles around 180°, perhaps in the helix 5/2 orientation. In contrast, the mCherry nanodiscs seem to adopt a more compact structure with mCherry-to-mCherry angles around 65° (Figure S9), perhaps in the helix 5/5 orientation. These changes may partially explain the larger sizes observed in SEC for sfGFP nanodiscs relative to mCherry nanodiscs. However, future research will be needed to test this hypothesis in more detail.

Another potential factor could be the conformational space of the fluorescent protein fusions. Even if the register is the same for all nanodiscs, each fluorescent protein may dynamically sample different conformations, which could alter the observed Stokes diameters. We expect that the fluorescent proteins are likely flexible and dynamic as they extend off the side of the nanodisc. However, further research will be needed to explore this in more detail.

A final potential factor in explaining these observations is that the SEC was calibrated on globular proteins and may not accurately reflect the size of more extended structures. SEC may also be biased by surface interactions with the column. Thus, the effect could simply be an artifact of the SEC measurement.

**Extended Discussion of CD-MS and MP Results**

Both control and fluorescent nanodiscs sometimes had 10–20% higher fill ratios than predicted based on the assembly recipes. Some degree of overfilling could be natural experimental variation. However, prior native MS data has consistently shown systematic shifts towards higher masses when using the gentlest conditions. For example, Keener *et al.* [15]found a fill of around 150 POPC molecules in a MSP1D1 nanodisc despite a predicted fill of 130 based on the 65:1 assembly ratio. Townsend *et al.* found a fill of 185 DMPC in MSP1D1 nanodiscs with a predicted fill of 160.[16] Certainly, there is flexibility in the amount of lipids a nanodisc can accommodate, but the consistent systematic shift also suggests that the molar absorptivity for the MSP proteins may be 10-20% off, leading to a lower protein concentration and higher relative amount of lipids than expected in the recipe. Prior literature has shown that deviations of this magnitude are common.[17]

Interestingly, we found that fluorescent nanodiscs generally produced better native mass spectra than conventional nanodiscs and seemed to be more stable in the mass spectrometer. Conventional nanodiscs were more likely to fragment into MSP-lipid complexes between 2000–6000 *m/z*. To ameliorate this effect, we had to add charge-reducing reagents,[15] which lowered the quality of the spectra. However, the fluorescent nanodiscs were generally more stable and better matched the gentler CD-MS data while still producing well resolved native mass spectra without any charge reducing agents. We hypothesize that the fluorescent proteins provide additional surface area to distribute charge over, which reduces the charge directly on the lipid bilayer. However, further research will be needed to explore this observation.

Prior literature[18] has also shown that mass photometry (MP) results are systematically low in mass for nanodiscs. The primary challenge is that lipids and proteins have different average refractive index values per unit mass. Thus, MP calibrations done on proteins tend to underestimate the mass of nanodiscs with lipid. Interestingly, our results seem to show more accurate measurements (defined by agreement with CD-MS results) for complexes with the protein fusions, likely because more of the mass is protein and thus better matches the calibration. Due to the different protein versus lipid content used here, we did not attempt to correct the results but report them as measured. However, it may be possible in future studies to use these sorts of constructs to systematically vary the lipid and protein content to create more accurate MP calibrations.

**Demonstration of Membrane Protein Incorporation**

We tested our fluorescent MSP constructs for membrane protein incorporation using *E. coli* aquaporin Z (AqpZ), which has previously been incorporated in nanodiscs.[16,19] The AqpZ construct has a GFP fusion, so we chose an sfCherry MSP to provide a different color to mark the MSP. Due to the size of AqpZ, we used an MSP1E3D1 construct with the N-terminal His-tag removed. We assembled the nanodiscs following established protocols[16,19] and purified the resulting mixture first by IMAC to isolate nanodiscs containing AqpZ and then by SEC. Empty nanodiscs from the same assembly reaction were collected from the IMAC flowthrough and also purified with SEC.

SDS-PAGE analysis revealed that the AqpZ nanodiscs had both red and green fluorescence and clear bands for the MSP and AqpZ (Figure S10A). Note, we did not boil the samples to avoid denaturing the fluorescent proteins, so AqpZ had a broad band and some dimer due to incomplete denaturation in SDS. Empty nanodiscs lacked the green fluorescence from the AqpZ bands. Imaging both the green and red channels confirmed that both proteins were present in the purified samples.

Next, we performed FRET on the purified AqpZ nanodiscs (Figure S10B), which showed higher FRET than empty nanodiscs from the same reaction. These data demonstrate that both proteins are in close proximity, consistent with them being co-incorporated inside the same nanodisc complex. Finally, MP analysis confirmed the AqpZ nanodiscs had the expected mass (Figure S10C). As discussed above, there can be systematic errors in MP results due to the lipids, but the high protein content of the AqpZ nanodiscs makes the MP measurement likely closer. The AqpZ tetramer has a mass of 208 kDa, and the MSP dimer has a mass of 111 kDa. Subtracting these from the measured MP mass of 451 kDa yields 132 kDa of lipid mass, for an estimated ratio of 195 lipids per nanodisc, which is consistent with prior native MS results[20] for AqpZ in POPC nanodiscs with MSP1E3D1.

These data demonstrate that fluorescent MSPs can be used to incorporate membrane proteins into nanodiscs. Furthermore, they demonstrate the value in having diverse fluorescent colors available. It was immediately clear by eye that the IMAC flowthrough was red and that the elution was brownish, indicating both red and green. It was also clear which fractions had the protein. Having two colors of fluorescence enables easy tracking of the membrane protein and MSP during nanodisc assembly.

**Supplemental Figures**

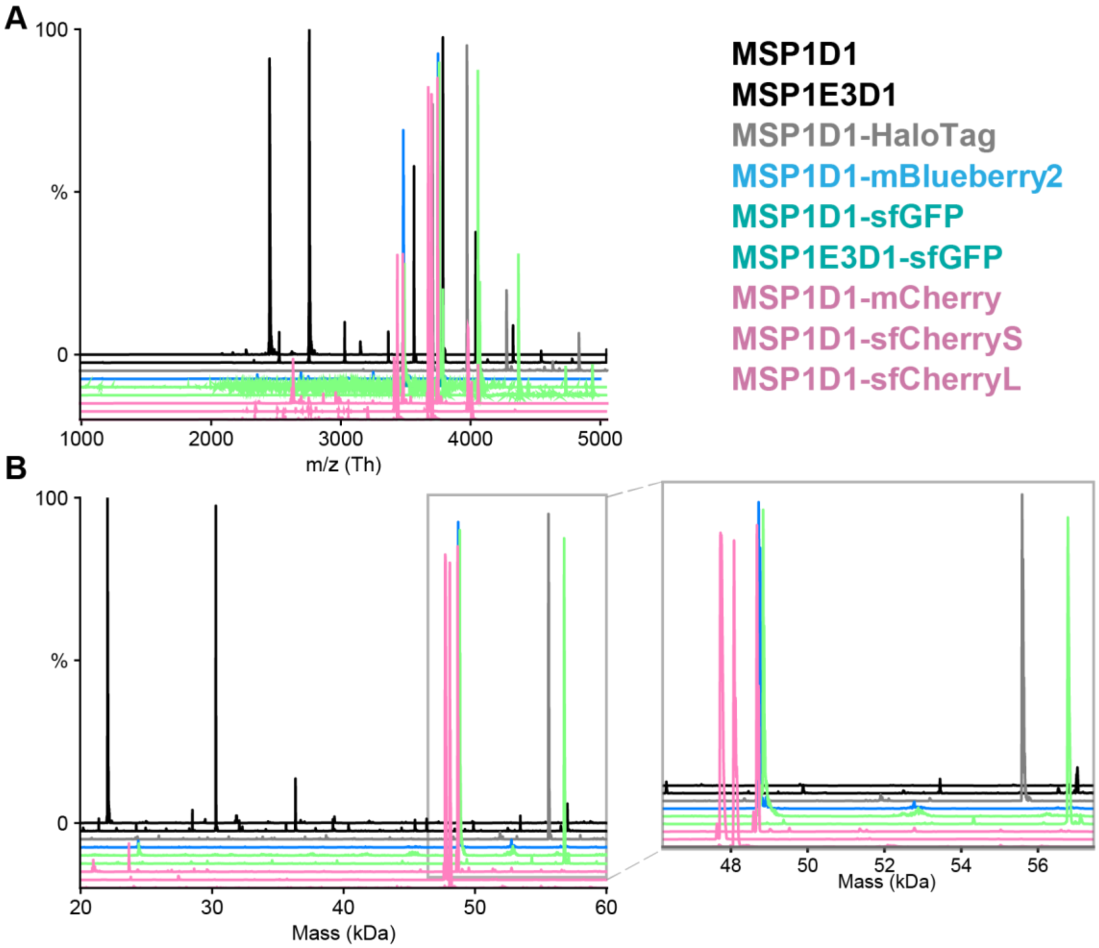

**Figure S1.** Representative mass spectrum for each free MSP with A) raw spectra of each MSP and B) deconvolved mass spectra for free MSP constructs with a zoomed in section at 45 to 60 kDa. Masses are provided in Table S4 below.

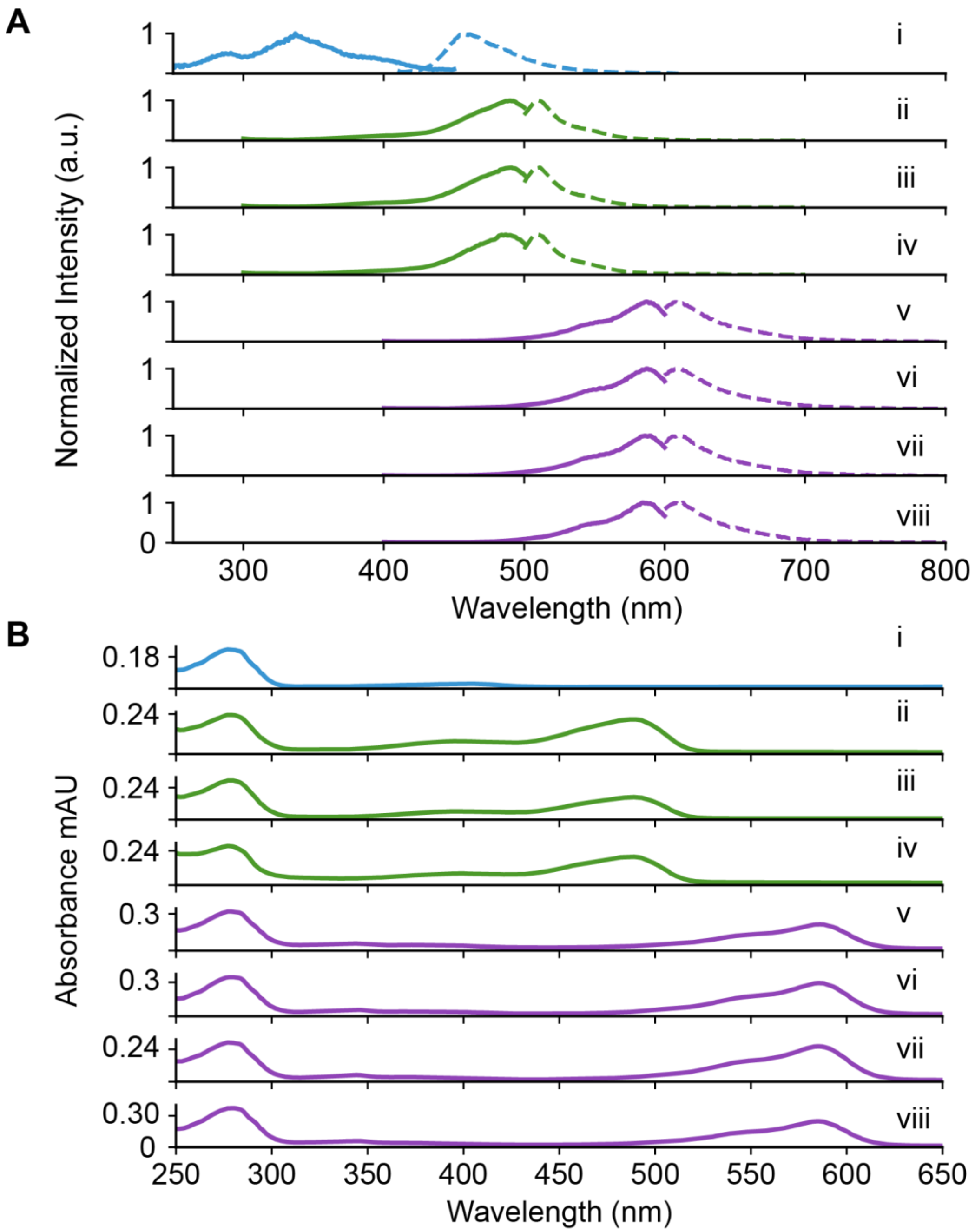

**Figure S2.** Fluorescence and absorbance spectra of fluorescent proteins. A) Fluorescent excitation (*solid*) and emission (*dashed*) spectra at 1 µM. B) UV-Vis spectra at a concentration of 5 µM, based on BCA analysis. Spectra are shown for: i) MSP1D1-mBlueberry2, ii) MSP1D1-sfGFP, iii) MSP1E3D1-sfGFP, iv) sfGFP-Annexin V, v) MSP1D1-mCherry, vi) MSP1D1-sfCherryS, vii) MSP1D1-sfCherryL, and viii) MSP1E3D1-sfCherry3C.

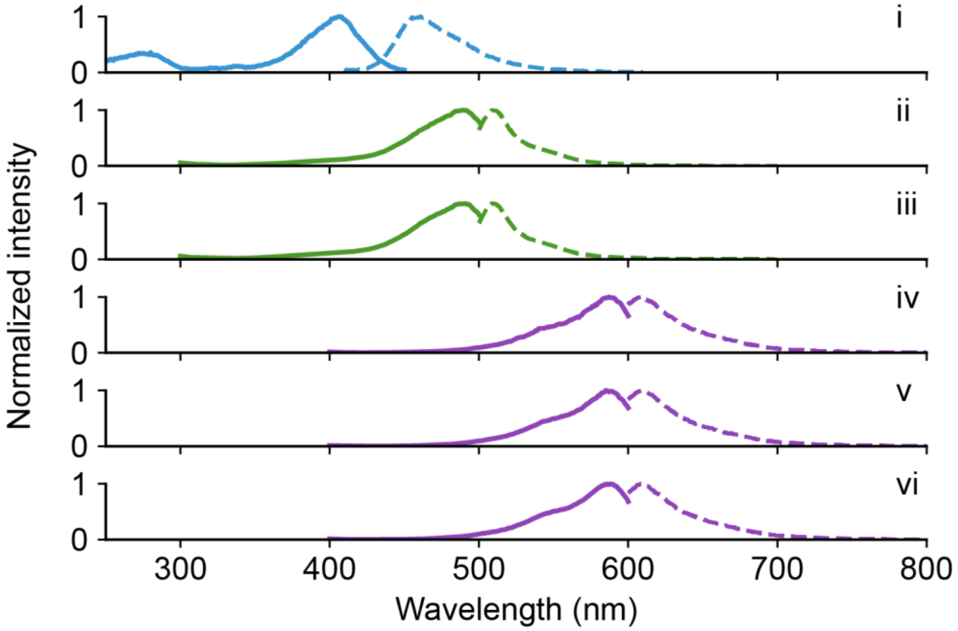

**Figure S3.** Nanodisc fluorescent excitation and emission spectra for nanodiscs formed with: i) MSP1D1-mBlueberry2, ii) MSP1D1-sfGFP, iii) MSP1E3D1-sfGFP, iv) MSP1D1-mCherry, v) MSP1D1-sfCherryS, and vi) MSP1D1-sfCherryL.

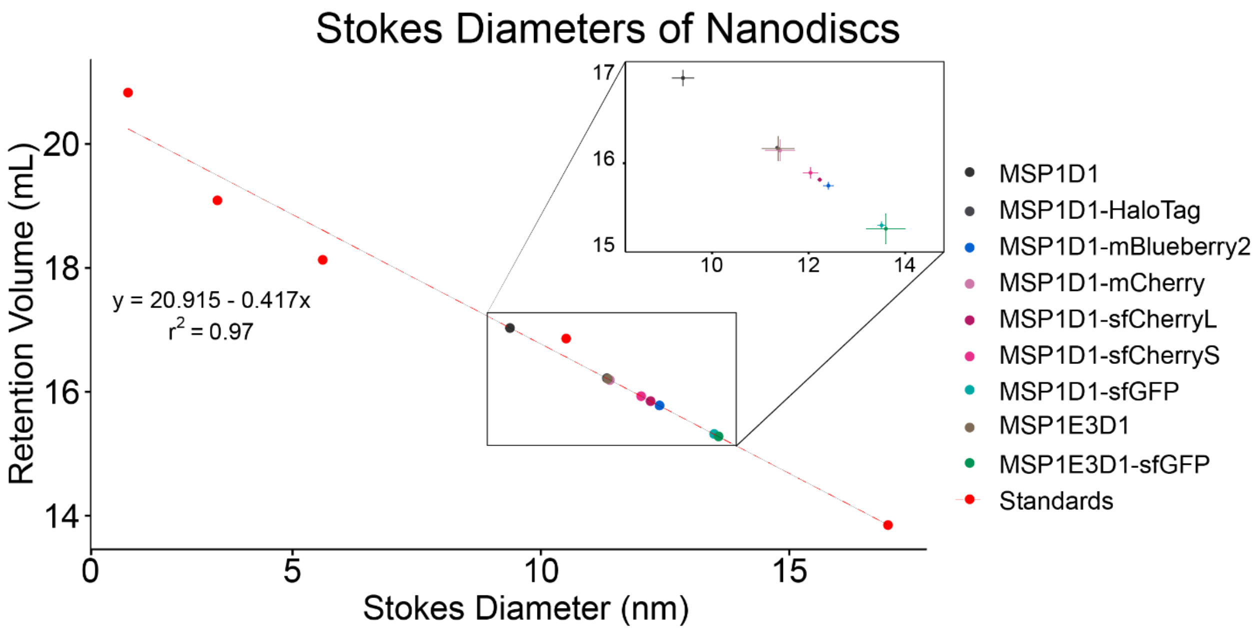

**Figure S4.** Stokes diameter of nanodiscs formed from different MSP belts as annotated. The zoomed region displays the nanodisc Stokes diameter. Error bars represent the standard deviation from triplicate measurements. Results are provided in Table S5 below.

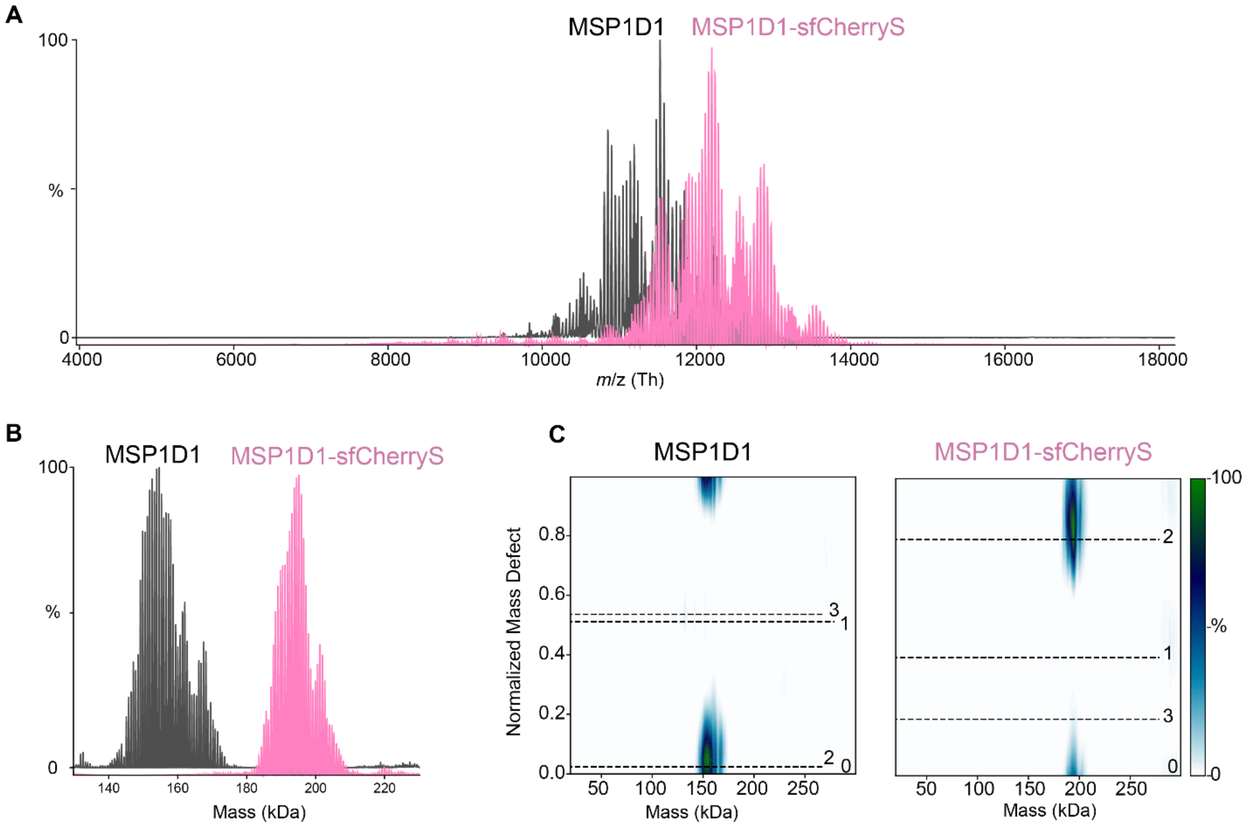

**Figure S5.** Native MS and macromolecular mass defect analysis of MSP1D1 nanodiscs and MSP1D1-sfCherryS nanodiscs. A) Raw mass spectra of control (*grey*) and MSP1D1-sfCherryS (*pink*) DMPC nanodiscs and B) deconvolved zero charge mass spectra of nanodiscs. The MSP1D1 mass distribution is centered around 152 kDa, and the MSP1D1-sfCherryS mass distribution is centered around 195 kDa. C) Normalized molecular mass defects for control nanodiscs (*left*) and MSP1D1-sfCherryS (*right*). Theoretical mass defects for different numbers of MSP belts (0, 1, 2, and 3) are annotated to the right on each plot. Both show only two MSP per nanodisc.

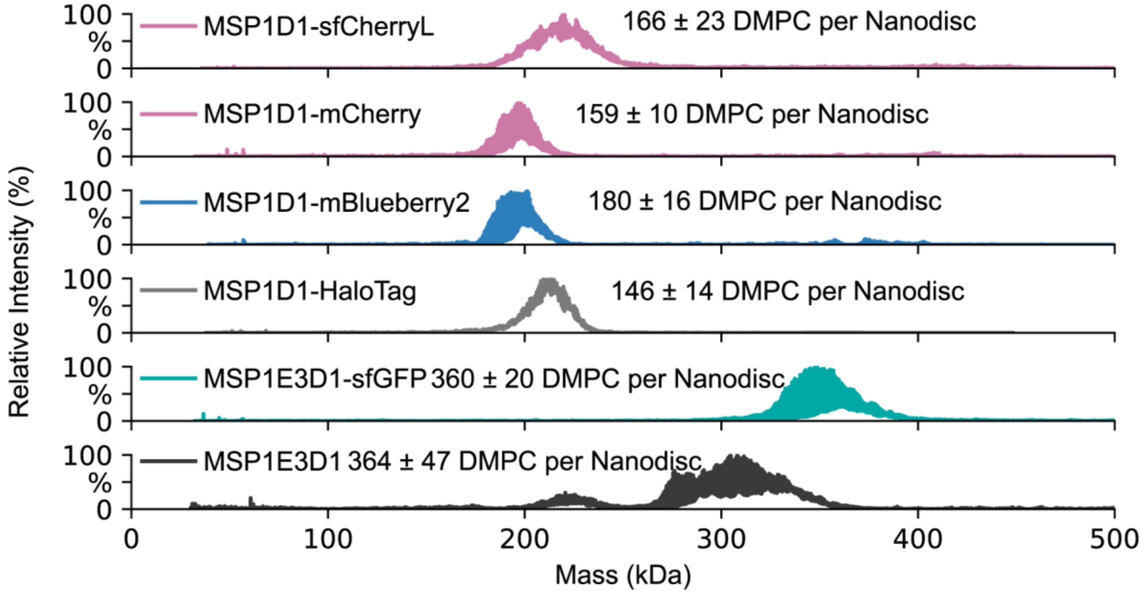

**Figure S6.** Representative CD-MS mass distributions of nanodiscs with additional MSP belts: A) MSP1D1-sfCherryL, B) MSP1D1-mCherry, C) MSP1D1-mBlueberry2, D) MSP1D1-HaloTag, and E) MSP1E3D1-sfGFP. The number of DMPC molecules per nanodisc was calculated by subtracting the mass of two MSP belts and dividing by the DMPC mass. Results are reported as mean and standard deviation of three replicate nanodisc assemblies. The MSP1E3D1 constructs are expected to have more lipids than MSP1D1-based constructs.[21]

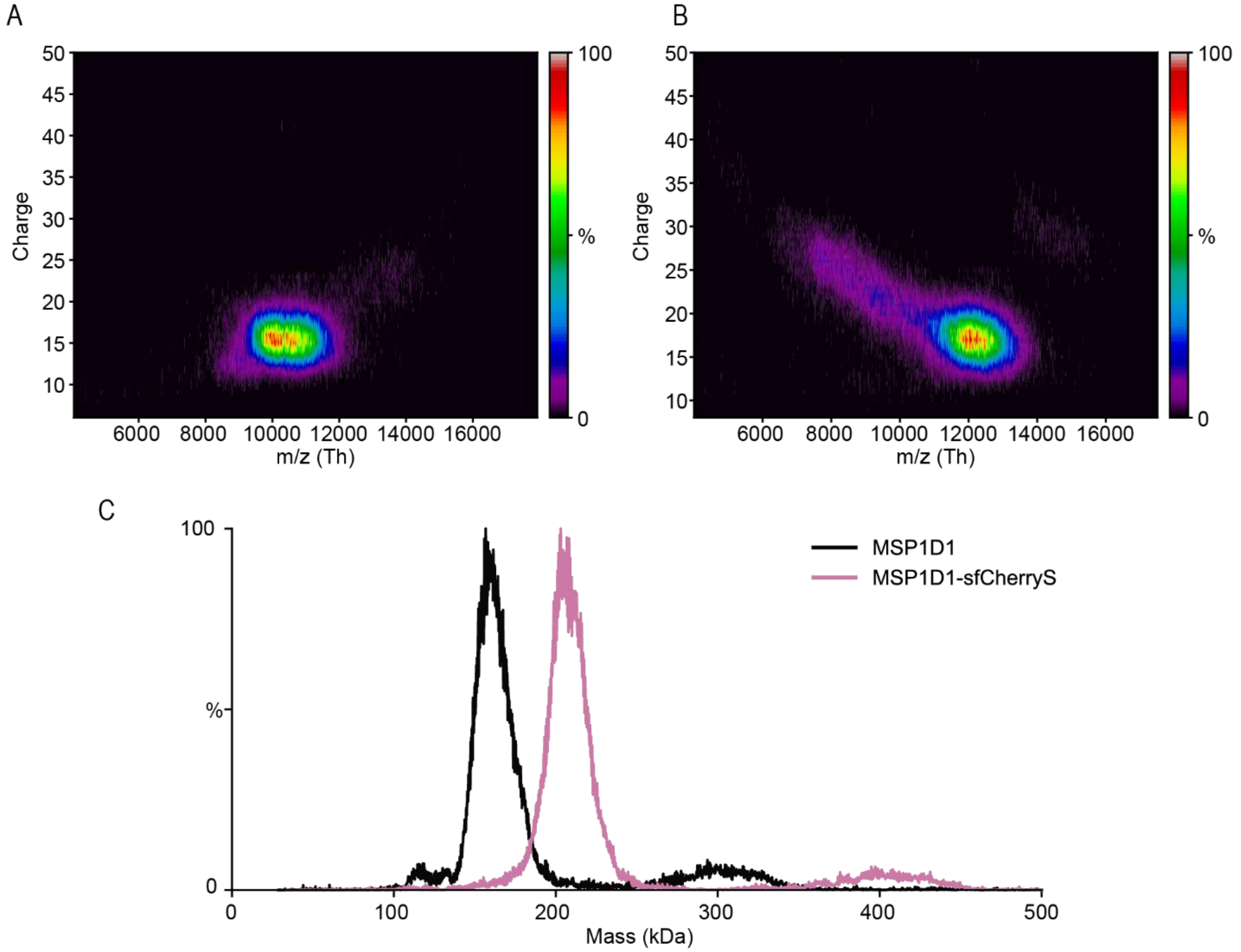

**Figure S7.** Raw CD-MS charge vs *m*/*z* data for A) MSP1D1 nanodiscs and B) MSP1D1-sfCherryS nanodiscs. C) Deconvolution shows predominantly single nanodisc complexes at the expected masses with a small amount of dimer at double the mass. Dimer peaks are observed at unique *m/z* values in A and B, demonstrating that these are not artifacts of CD-MS analysis.

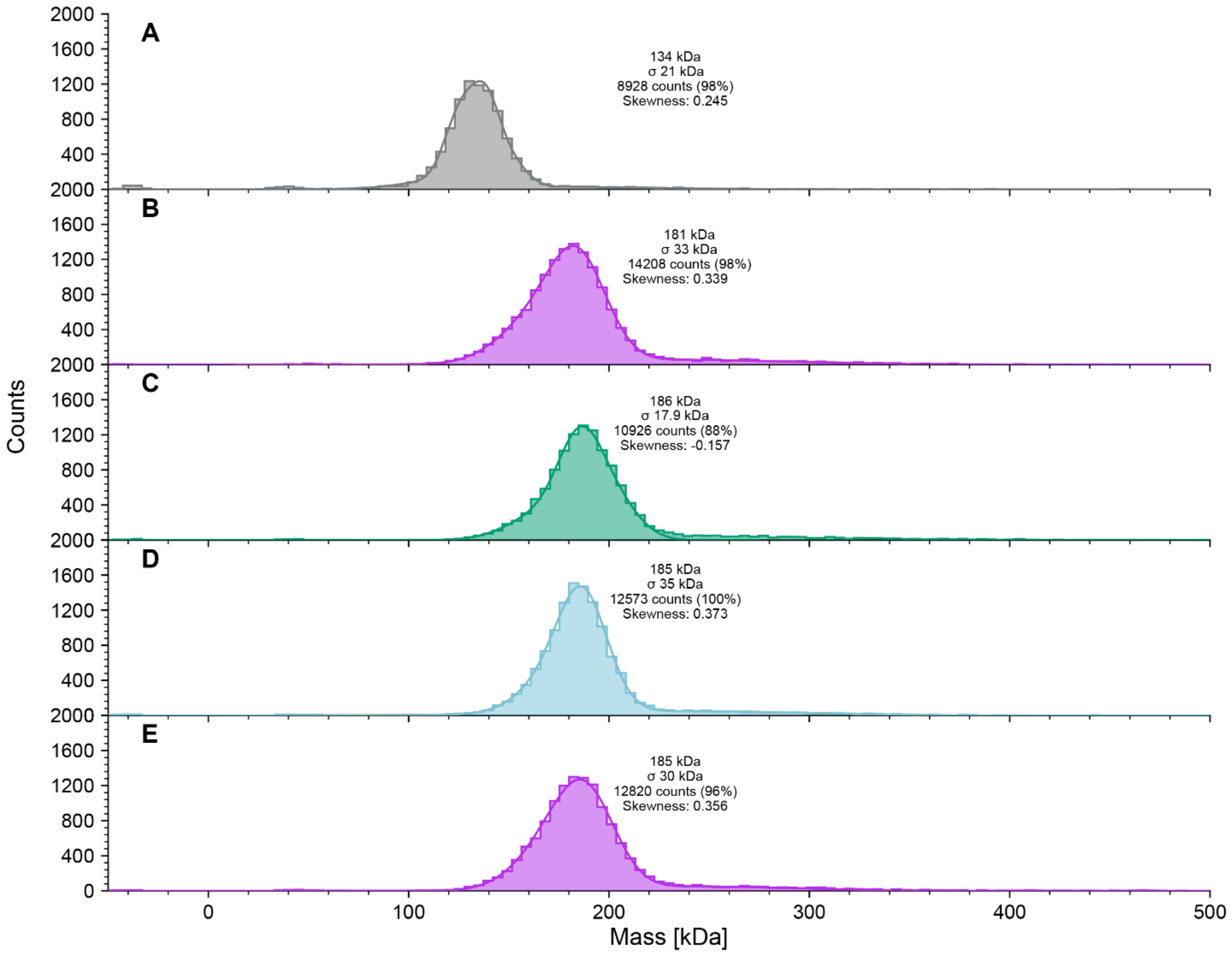

**Figure S8.** Representative mass photometry data for triplicate DMPC nanodiscs formed with: A) MSP1D1. B) MSP1D1-mCherry, C) MSP1D1-sfGFP, D) MSP1D1-mBluebery2, and E) MSP1D1-sfCherryS. No significant fraction of dimers is observed. Results are provide in Table S6 below.

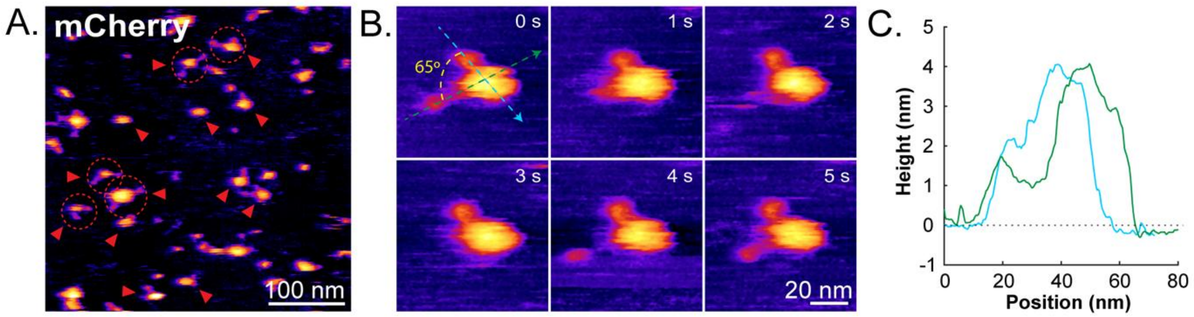

**Figure S9.** Structural characterization of mCherry nanodiscs. (A) Representative HS-AFM images of the mCherry nanodiscs deposited on mica substrate. Red arrows indicate isolated nanodiscs. Dashed circles highlight the nanodiscs in which two fused-mCherry molecules are visible. (B) Real-time HS-AFM images showing the structural dynamics of an individual mCherry nanodisc. The two fused mCherry molecules show an apparent orientation angle of approximately 65°. (C) Cross-sectional height profiles of the mCherry nanodisc shown in (B) at 0 s.

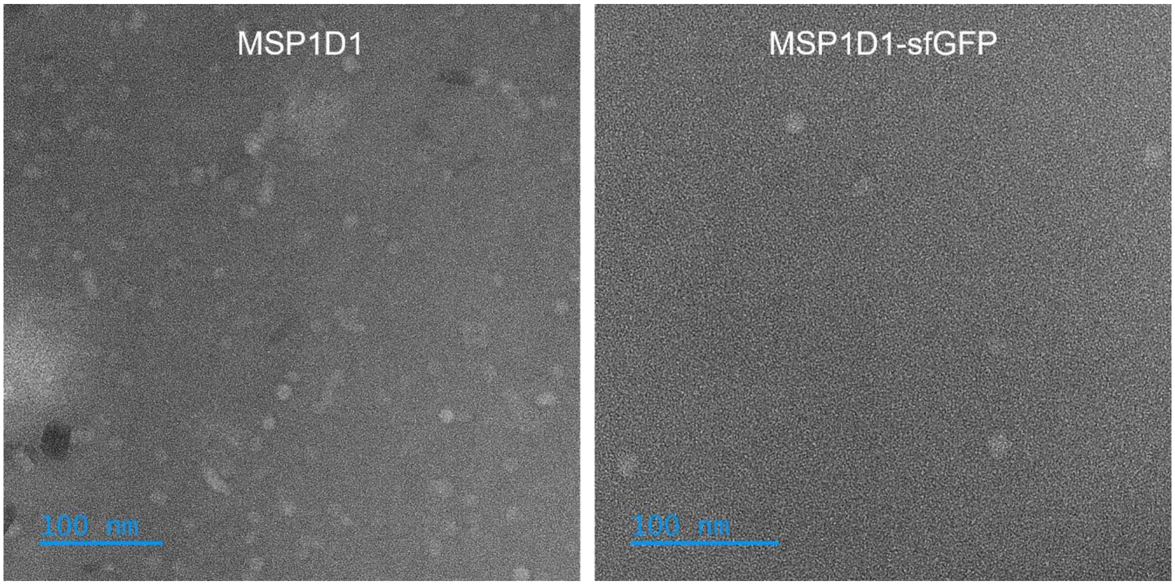

**Figure S10.** Negative stain TEM images of MSP1D1 (*left*) and MSP1D1-sfGFP nanodiscs (*right*). Samples were stained with uranyl acetate on a carbon grid and imaged on a JEOL 1400 S/TEM. For 4 particles in each, the ImageJ line tool was used to measure the particle diameters. MSP1D1 nanodiscs had an average diameter of 10.4 ± 0.4 nm and MSP1D1-sfGFP nanodiscs had an average diameter of 16.8 ± 1.9 nm.

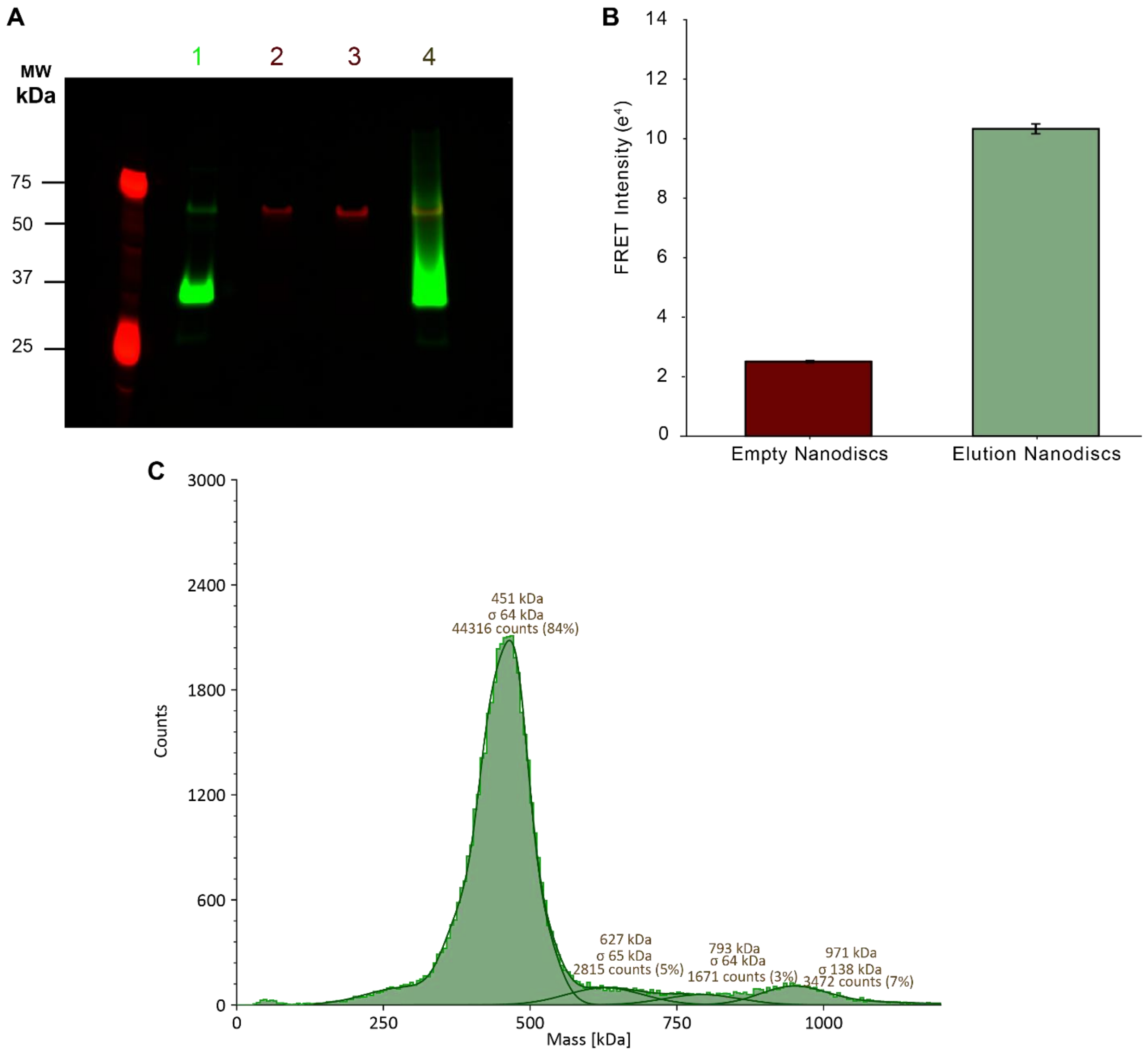

**Figure S11.** Characterization of AqpZ nanodiscs. A) In-gel fluorescent image of SDS-PAGE in both red and green colors. Lanes contained: 1) AqpZ in detergent, 2) free MSP1E3D1-sfCherry3C, 3) empty nanodiscs from the flow-through of the IMAC column that lacked the AqpZ, and 4) purified AqpZ nanodiscs. B) FRET intensities of the empty nanodiscs compared to AqpZ nanodiscs. Error bars report standard deviations on triplicate measurements of a single assembly reaction. C) Mass photometry of the AqpZ nanodiscs.

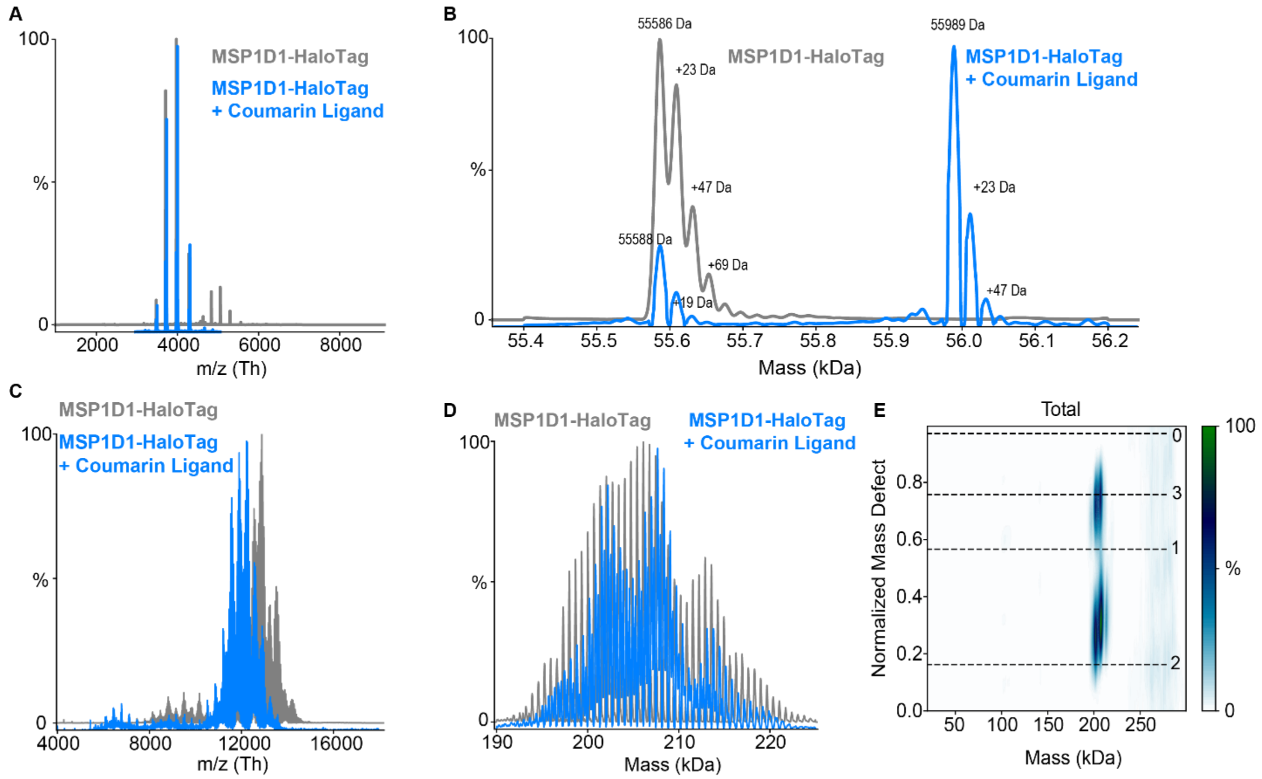

**Figure S12.** MSP1D1-HaloTag labeling with HaloTag-coumarin label. A) Raw native mass spectra of MSP1D1-HaloTag unlabeled (*grey*) and labeled with HaloTag Coumarin ligand (*blue*) overlaid. B) Deconvolved mass show around 70% labeling. C) Raw mass spectra of MSP1D1-HaloTag unlabeled and labeled nanodiscs overlaid. D) Deconvolved mass distribution showing that labeling did not perturb the overall mass distribution. E) Mass defect analysis for labeled nanodiscs showing that the nanodiscs had 1–2 labels per nanodiscs. Mass defect peaks often run slightly higher in mass than predicted due to adduction, as seen in B.

### Supplementary Tables

**Table S1.** Fluoro-MSP sequences with references, Addgene links, and Addgene plasmid numbers. Constructs are colored with the His-tag in *red*; post-his linker in *black*; TEV site in *grey*; MSP in *blue*; post-MSP linker in *black*; GFP in *green*, mCherry, sfCherry, and sfCherry3C in *purple*, mBlueberry2 in *light blue*, HaloTag in *dark blue,* and Annexin V in *black.*

| Construct Name | Sequences | References |
|---|---|---|
| **pMSP1D1_sfGFP** | MGHHHHHHHDYDIPTTENLYFQ\|GSTFSKLREQLGPVTQEFWDNLEKETEGLRQEMSKDLEEVKAKVQPYLDDFQKKWQEEMELYRQKVEPLRAELQEGARQKLHELQEKLSPLGEEMRDRARAHVDALRTHLAPYSDELRQRLAARLEALKENGGARLAEYHAKATEHLSTLSEKAKPALEDLRQGLLPVLESFKVSFLSALEEYTKKLNTQQGSKGEELFTGVVPILVELDGDVNGHKFSVRGEGEGDATNGKLTLKFICTTGKLPVPWPTLVTTLTYGVQCFSRYPDHMKRHDFFKSAMPEGYVQERTISFKDDGTYKTRAEVKFEGDTLVNRIELKGIDFKEDGNILGHKLEYNFNSHNVYITADKQKNGIKANFKIRHNVEDGSVQLADHYQQNTPIGDGPVLLPDNHYLSTQSVLSKDPNEKRDHMVLLEFVTAAGITHGMDELYK | A portion of this plasmid was derived from a plasmid made by Stephen Sligar (https://www.addgene.org/20061/), and the fluorescent protein sequence was derived from https://www.fpbase.org/protein/superfolder-gfp/ (DOI: 10.1038/nbt1172).<br>Plasmid link: https://www.addgene.org/Michael_Marty/<br>Plasmid number: 248920 |
| **pMSP1D1_mCherry** | MGHHHHHHHDYDIPTTENLYFQGSTFSKLREQLGPVTQEFWDNLEKETEGLRQEMSKDLEEVKAKVQPYLDDFQKKWQEEMELYRQKVEPLRAELQEGARQKLHELQEKLSPLGEEMRDRARAHVDALRTHLAPYSDELRQRLAARLEALKENGGARLAEYHAKATEHLSTLSEKAKPALEDLRQGLLPVLESFKVSFLSALEEYTKKLNTQQGSKGEEDNMAIIKEFMRFKVHMEGSVNGHEFEIEGEGEGRPYEGTQTAKLKVTKGGPLPFAWDILSPQFMYGSKAYVKHPADIPDYLKLSFPEGFKWERVMNFEDGGVVTVTQDSSLQDGEFIYKVKLRGTNFPSDGPVMQKKTMGWEASSERMYPEDGALKGEIKQRLKLKDGGHYDAEVKTTYKAKKPVQLPGAYNVNIKLDITSHNEDYTIVEQYERAEGRHSTGGMDELYK | A portion of this plasmid was derived from a plasmid made by Stephen Sligar (https://www.addgene.org/20061/), and the fluorescent protein sequence was derived from https://www.fpbase.org/protein/mcherry/ (DOI: 10.1038/nbt1037).<br>Plasmid link: https://www.addgene.org/Michael_Marty/<br>Plasmid number: 248922 |
| **pMSP1D1_sfCherryS** | MGHHHHHHHDYDIPTTENLYFQGSTFSKLREQLGPVTQEFWDNLEKETEGLRQEMSKDLEEVKAKVQPYLDDFQKKWQEEMELYR | A portion of this plasmid was derived from a plasmid made by Stephen Sligar |

| | | |
|---|---|---|
| | QKVEPLRAELQEGARQKLHELQEKLSPLGEEMRDRARAHVDALRTHLAPYSDELRQRLAARLEALKENGGARLAEYHAKATEHLSTLSEKAKPALEDLRQGLLPVLESFKVSFLSALEEYTKKLNTQQGSEEDNNMAIIKEFMRFKVHMEGSVNGHEFEIEGEGEGHPYEGTQTAKLKVTKGGPLPFAWDILSPQFMYGSKAYVKHPADIPDYLKLSFPEGFTWERVMNFEDGGVVTVTQDSSLQDGEFIYKVKLLGTNFPSDGPVMQKKTNGWEASTERMYPEDGALKGEINQRLKLKDGGHYDAEVKTTYKAKKPVQLPGAYNVDIKLDITSHNEDYTIVEQYERAEGRHSTGG | (https://www.addgene.org/20061/), and the fluorescent protein sequence was derived from https://www.fpbase.org/protein/sfcherry/ (DOI: 10.1107/s0907444913024608). Plasmid link: https://www.addgene.org/Michael_Marty/ Plasmid number: 248923 |
| **pMSP1D1_sfCherryL** | MGHHHHHHHDYDIPTTENLYFQGSTFSKLREQLGPVTQEFWDNLEKETEGLRQEMSKDLEEVKAKVQPYLDDFQKKWQEEMELYRQKVEPLRAELQEGARQKLHELQEKLSPLGEEMRDRARAHVDALRTHLAPYSDELRQRLAARLEALKENGGARLAEYHAKATEHLSTLSEKAKPALEDLRQGLLPVLESFKVSFLSALEEYTKKLNTQGGGGSGGGGSEEDNNMAIIKEFMRFKVHMEGSVNGHEFEIEGEGEGHPYEGTQTAKLKVTKGGPLPFAWDILSPQFMYGSKAYVKHPADIPDYLKLSFPEGFTWERVMNFEDGGVVTVTQDSSLQDGEFIYKVKLLGTNFPSDGPVMQKKTNGWEASTERMYPEDGALKGEINQRLKLKDGGHYDAEVKTTYKAKKPVQLPGAYNVDIKLDITSHNEDYTIVEQYERAEGRHSTGG | A portion of this plasmid was derived from a plasmid made by Stephen Sligar (https://www.addgene.org/20061/), and the fluorescent protein sequence was derived from https://www.fpbase.org/protein/sfcherry/ (DOI: 10.1107/s0907444913024608). Plasmid link: https://www.addgene.org/Michael_Marty/ Plasmid number: 248924 |
| **pMSP1D1_mBlueberry2** | MGHHHHHHHDYDIPTTENLYFQGSTFSKLREQLGPVTQEFWDNLEKETEGLRQEMSKDLEEVKAKVQPYLDDFQKKWQEEMELYRQKVEPLRAELQEGARQKLHELQEKLSPLGEEMRDRARAHVDALRTHLAPYSDELRQRLAARLEALKENGGARLAEYHAKATEHLSTLSEKAKPALEDLRQGLLPVLESFKVSFLSALEEYTKKLNTQQGSKGEENNVAIIKEFMRFKVHMEGSVNGHEFEIEGEGEGRPYEGTQTAKLKVTKGGPLPFAWDILSPQFLFGSKVYIKHPADIPDYFKLSFPEGFKWERVMNFEDGGVVTVTQDSSLQDGVFIYKVKLRGTNFPSDGPVMQKKTMGWEAFSERMYPEDGALKSEIKTRLKLKDGGHYDAEVKTTYKAKKPVQLPGAYNVNIKLDIVSHNEDYTIVEQYERAEGRHSTGGMDELYK | A portion of this plasmid was derived from a plasmid made by Stephen Sligar (https://www.addgene.org/20061/), and the fluorescent protein sequence was derived from https://www.fpbase.org/protein/mblueberry2/ (DOI: 10.1021/bi700199g). Plasmid link: https://www.addgene.org/Michael_Marty/ Plasmid number: 248925 |
| **pMSP1D1_HaloTag** | MGHHHHHHHDYDIPTTENLYFQGSTFSKLREQLGPVTQEFWDNLEKETEGLRQEMSKDLEEVKAKVQPYLDDFQKKWQEEMELYRQKVEPLRAELQEGARQKLHELQEKLSPL | A portion of this plasmid was derived from a plasmid made by Stephen Sligar (https://www.addgene.org/20061/), and the HaloTag protein sequence was |

| | | |
|---|---|---|
| | GEEMRDRARAHVDALRTHLAPYSDELRQ<br>RLAARLEALKENGGARLAEYHAKATEHLS<br>TLSEKAKPALEDLRQGLLPVLESFKVSFLS<br>ALEEYTKKLNTQQGSIGTGFPFDPHYVEV<br>LGERMHYVDVGPRDGTPVLFLHGNPTSS<br>YVWRNIIPHVAPTHRCIAPDLIGMGKSDKP<br>DLGYFFDDHVRFMDAFIEALGLEEVVLVIH<br>DWGSALGFHWAKRNPERVKGIAFMEFIR<br>PIPTWDEWPEFARETFQAFRTTDVGRKLII<br>DHNVFIEGTLRMGVVRPLTEVEMDHYRE<br>PFLNPVDREPLWRFPNELPIAGEPANIVAL<br>VEEYMDWLHQSPVPKLLFWGTPGVLIPP<br>AEAARLAKSLPNCKAVDIGPGLNLLQEDN<br>PDLIGSEIARWLSTLEIS | derived from (DOI: https://doi.org/10.1038/s41592-021-01341-x).<br>Plasmid link: https://www.addgene.org/Michael_Marty/<br>Plasmid number: 248926 |
| **pMSP1E3D1_sfGFP** | MGHHHHHHHDYDIPTTENLYFQGSTFSKL<br>REQLGPVTQEFWDNLEKETEGLRQEMSK<br>DLEEVKAKVQPYLDDFQKKWQEEMELYR<br>QKVEPLRAELQEGARQKLHELQEKLSPL<br>GEEMRDRARAHVDALRTHLAPYLDDFQK<br>KWQEEMELYRQKVEPLRAELQEGARQKL<br>HELQEKLSPLGEEMRDRARAHVDALRTH<br>LAPYSDELRQRLAARLEALKENGGARLAE<br>YHAKATEHLSTLSEKAKPALEDLRQGLLP<br>VLESFKVSFLSALEEYTKKLNTQQGSKGE<br>ELFTGVVPILVELDGDVNGHKFSVRGEGE<br>GDATNGKLTLKFICTTGKLPVPWPTLVTTL<br>TYGVQCFSRYPDHMKRHDFFKSAMPEG<br>YVQERTISFKDDGTYKTRAEVKFEGDTLV<br>NRIELKGIDFKEDGNILGHKLEYNFNSHNV<br>YITADKQKNGIKANFKIRHNVEDGSVQLAD<br>HYQQNTPIGDGPVLLPDNHYLSTQSVLSK<br>DPNEKRDHMVLLEFVTAAGITHGMDELYK | A portion of this plasmid was derived from a plasmid made by Stephen Sligar (https://www.addgene.org/20066/), and the fluorescent protein sequence was derived from https://www.fpbase.org/protein/superfolder-gfp/ (DOI: 10.1038/nbt1172).<br>Plasmid link: https://www.addgene.org/Michael_Marty/<br>Plasmid number: 248928 |
| **pMSP1E3D1_mCherry** | MGHHHHHHHDYDIPTTENLYFQGSTFSKL<br>REQLGPVTQEFWDNLEKETEGLRQEMSK<br>DLEEVKAKVQPYLDDFQKKWQEEMELYR<br>QKVEPLRAELQEGARQKLHELQEKLSPL<br>GEEMRDRARAHVDALRTHLAPYLDDFQK<br>KWQEEMELYRQKVEPLRAELQEGARQKL<br>HELQEKLSPLGEEMRDRARAHVDALRTH<br>LAPYSDELRQRLAARLEALKENGGARLAE<br>YHAKATEHLSTLSEKAKPALEDLRQGLLP<br>VLESFKVSFLSALEEYTKKLNTQQGSKGE<br>EDNMAIIKEFMRFKVHMEGSVNGHEFEIE<br>GEGEGRPYEGTQTAKLKVTKGGPLPFAW<br>DILSPQFMYGSKAYVKHPADIPDYLKLSFP<br>EGFKWERVMNFEDGGVVTVTQDSSLQD<br>GEFIYKVKLRGTNFPSDGPVMQKKTMGW<br>EASSERMYPEDGALKGEIKQRLKLKDGG | A portion of this plasmid was derived from a plasmid made by Stephen Sligar (https://www.addgene.org/20066/), and the fluorescent protein sequence was derived from https://www.fpbase.org/protein/mcherry/ (DOI: 10.1038/nbt1037).<br>Plasmid link: https://www.addgene.org/Michael_Marty/<br>Plasmid number: 248929 |

| | | |
|---|---|---|
| | HYDAEVKTTYKAKKPVQLPGAYNVNIKLDI<br>TSHNEDYTIVEQYERAEGRHSTGGMDEL<br>YK | |
| **pMSP1E3D1_mBlueberry2** | MGHHHHHHHDYDIPTTENLYFQGSTFSKL<br>REQLGPVTQEFWDNLEKETEGLRQEMSK<br>DLEEVKAKVQPYLDDFQKKWQEEMELYR<br>QKVEPLRAELQEGARQKLHELQEKLSPL<br>GEEMRDRARAHVDALRTHLAPYLDDFQK<br>KWQEEMELYRQKVEPLRAELQEGARQKL<br>HELQEKLSPLGEEMRDRARAHVDALRTH<br>LAPYSDELRQRLAARLEALKENGGARLAE<br>YHAKATEHLSTLSEKAKPALEDLRQGLLP<br>VLESFKVSFLSALEEYTKKLNTQQGSKGE<br>ENNVAIIKEFMRFKVHMEGSVNGHEFEIE<br>GEGEGRPYEGTQTAKLKVTKGGPLPFAW<br>DILSPQFLFGSKVYIKHPADIPDYFKLSFPE<br>GFKWERVMNFEDGGVVTVTQDSSLQDG<br>VFIYKVKLRGTNFPSDGPVMQKKTMGWE<br>AFSERMYPEDGALKSEIKTRLKLKDGGHY<br>DAEVKTTYKAKKPVQLPGAYNVNIKLDIVS<br>HNEDYTIVEQYERAEGRHSTGGMDELYK | A portion of this plasmid was derived from a plasmid made by Stephen Sligar (https://www.addgene.org/20066/), and the fluorescent protein sequence was derived from https://www.fpbase.org/protein/mblueberry2/ (DOI: 10.1021/bi700199g).<br>Plasmid link: https://www.addgene.org/Michael_Marty/<br>Plasmid number: 248930 |
| **pMSP1E3D1_sfCherry3C** | MGHHHHHHHDYDIPTTENLYFQGSTFSKL<br>REQLGPVTQEFWDNLEKETEGLRQEMSK<br>DLEEVKAKVQPYLDDFQKKWQEEMELYR<br>QKVEPLRAELQEGARQKLHELQEKLSPL<br>GEEMRDRARAHVDALRTHLAPYLDDFQK<br>KWQEEMELYRQKVEPLRAELQEGARQKL<br>HELQEKLSPLGEEMRDRARAHVDALRTH<br>LAPYSDELRQRLAARLEALKENGGARLAE<br>YHAKATEHLSTLSEKAKPALEDLRQGLLP<br>VLESFKVSFLSALEEYTKKLNTQQGSEED<br>NMAIIKEFMRFKVHMEGSVNGHEFEI EGE<br>GEGHPYEGTQTARLKVTKGDPLPFAWDIL<br>SPQFMYGSKAYVKHPADIPDYLKLSFPEG<br>FTWERVMNFEDGGVVAVTQDSSLQDGQ<br>FIYKVKLLGINFPSDGPVMQKKTMGWEAS<br>TERMYPEDGALKGEINQRLKLKDGGHYD<br>AEVKTTYRAKKPVQLPGAYDVDIKLDITSH<br>NEDYTIVEQYERAEARHST | A portion of this plasmid was derived from a plasmid made by Stephen Sligar (https://www.addgene.org/20066/), and the fluorescent protein sequence was derived from https://www.fpbase.org/protein/sfcherry3c/ ( DOI: 10.1038/s42003-019-0589-x.)<br>Plasmid number: 257989 |

| | | |
|---|---|---|
| **AnnexinV_sfGFP** | MGHHHHHHHDYDIPTTSKGEELFTGVVPI<br>LVELDGDVNGHKFSVRGEGEGDATNG<br>KLTLKFICTTGKLPVPWPTLVTTLTYGVQC<br>FSRYPDHMKR<br>HDFFKSAMPEGYVQERTISFKDDGTYKTR<br>AEVKFEGDTLVNRIELKGIDF<br>KEDGNILGHKLEYNFNSHNVYITADKQKN<br>G<br>IKANFKIRHNVEDGSVQLADHYQQNTPIG<br>DGPVLLPDNHYLSTQSVLSKDPNEKRDH<br>MVLLEFVTAAGITHGMDELYKENLYFQGS<br>MAQVLRGTVTDFPGFDERADAETLRKAM<br>KGLGTDEESILTLLTSRSNAQRQEISAAFK<br>TLFGRDLLDDLKSELTGKFEKLIVALMKPS<br>RLYDAYELKHALKGAGTNEKVLTEIIASRT<br>PEELRAIKQVYEEEYGSSLEDDVVGDTSG<br>YYQRMLVVLLQANRDPDAGIDEAQVEQD<br>AQALFQAGELKWGTDEEKFITIFGTRSVS<br>HLRKVFDKYMTISGFQIEETIDRETSGNLE<br>QLLLAVVKSIRSIPAYLAETLYYAMKGAGT<br>DDHTLIRVMVSRSEIDLFNIRKEFRKNFAT<br>SLYSMIKGDTSGDYKKALLLLCGEDD | The fluorescent protein sequence was derived from https://doi.org/10.3389/fphys.2017.00317<br>Annexin V UniProt Accession: P08758<br>Plasmid number: 257992 |

**Table S2.** Selected native mass spectrometry settings for purified MSP protein analysis.

| Relevant Settings | Value |
|---|---|
| **Polarity** | Positive |
| **Maximum Inject Time** | 20 |
| **Spray voltage (kV)** | 1.1–1.3 |
| **Microscans** | 10 |
| **Capillary temperature (°C)** | 200 |
| **In-source trapping** | On |
| **Desolvation voltage (V)** | 0–150 |
| **Desolvation timing** | 10 |
| **Trapping Voltage (V)** | 40 |
| **HCD** | -35 |
| **In-source CID (eV)** | 50 |
| **Injection flatopole (V)** | 5.0 |
| **Bent flatopole (V)** | 2.0 |
| **Transfer Multipole** | 0 |
| **Trapping Gas Pressure** | 7.0 |
| **AGC Target** | 2e4 |
| **S-lens RF level** | 200 |
| **Scan Range** | 1,000–9,000 |
| **Resolution (at 200 m/z)** | 15,000–240,000 |

**Table S3.** Selected mass spectrometry settings for native and CD-MS for nanodiscs. Settings that did not change between the two tune files are denoted with (-)

| Relevant Settings | Native MS | CD-MS |
|---|---|---|
| **Spray voltage (kV)** | 1.1–1.3 | - |
| **Microscans** | 10 | 1 |
| **Capillary temperature (°C)** | 200 | - |
| **In-source trapping** | On | - |
| **Desolvation voltage (V)** | -150V | - |
| **Desolvation timing** | 10 | 4 |
| **Injection flatopole (V)** | 5.0 | - |
| **Bent flatopole (V)** | 2.0 | - |
| **Transfer Multipole** | 0 | - |
| **Trapping Gas Pressure** | 7.0 | 0.5 |
| **S-lens RF level** | 0.0 | - |
| **DMT** | Off | On |
| **Resolution (at 200 m/z)** | 7,500 | 240,000 |

**Table S4.** Predicted and measured masses of MSP constructs. Theoretical masses were calculated with Expasy ProtParam using the construct sequence as input and considering only the protein mass, not the expected mass change from the fluorophore formation. We elected not to include the fluorophore covalent mass changes in the theoretical masses because they are not known in all cases. Similarly, the MSP1D1-HaloTag-Coumarin theoretical mass was calculated as the protein sequence mass plus the theoretical mass of HaloTag-Coumarin ligand, without factoring in the loss of Cl during covalent attachment. Mass differences were measured as theoretical mass – measured mass.

| Protein | Theoretical Polypeptide Mass (Da) | Measured Mass (Da) | Mass Difference (Da) |
|---|---|---|---|
| **MSP1D1** | 22043 | 22043 | 0 |
| **MSP1E3D1** | 30285 | 30285 | 0 |
| **MSP1D1-sfGFP** | 48861 | 48840 | 21 |
| **MSP1E3D1-sfGFP** | 56799 | 56779 | 20 |
| **MSP1D1-sfCherryS** | 47746 | 47729 | 17 |
| **MSP1D1-sfCherryL** | 48104 | 48082 | 22 |
| **MSP1D1-mCherry** | 48702 | 48680 | 22 |
| **MSP1D1-mBlueberry2** | 48743 | 48724 | 19 |
| **MSP1D1-sfCFP** | 48884 | 48864 | 20 |
| **MSP1D1-HaloTag** | 55587 | 55586 | 1 |
| **MSP1D1-HaloTag-Coumarin** | 56026 | 55989 | 37 |

**Table S5.** Calculated Stokes diameters for each nanodisc. Retention volumes were collected for triplicate assemblies and averaged. Stokes diameters were calculated by linear regression using SEC calibrants of known diameter.

| Nanodisc MSP | Average Retention Volume (mL) | Standard Deviation (mL) | Average Stokes Diameter (nm) | Standard Deviation (nm) |
|---|---|---|---|---|
| **MSP1D1** | 16.99 | 0.09 | 9.32 | 0.22 |
| **MSP1E3D1** | 16.17 | 0.14 | 11.36 | 0.33 |
| **MSP1D1-sfGFP** | 15.28 | 0.04 | 13.58 | 0.08 |
| **MSP1E3D1-sfGFP** | 15.24 | 0.17 | 14.04 | 0.40 |
| **MSP1D1-sfCherry Short** | 15.89 | 0.06 | 12.15 | 0.15 |
| **MSP1D1-sfCherry Long** | 15.81 | 0.02 | 12.24 | 0.04 |
| **MSP1D1-mCherry** | 16.15 | 0.12 | 11.24 | 0.30 |
| **MSP1D1-mBlueberry2** | 15.74 | 0.04 | 12.48 | 0.10 |
| **MSP1D1-Halotag** | 16.18 | 0.01 | 11.36 | 0.01 |

**Table S6.** Average masses and standard deviations from triplicate nanodisc assemblies from mass photometry measurements.

| Nanodisc MSP | Average Mass (kDa) |
|---|---|
| **MSP1D1** | 137 ± 5 |
| **MSP1D1-sfGFP** | 190 ± 8 |
| **MSP1D1-sfCherryS** | 189 ± 6 |
| **MSP1D1-mCherry** | 184 ± 6 |
| **MSP1D1-mBlueberry2** | 189 ± 6 |
